# Cellular profiling of a recently-evolved social behavior

**DOI:** 10.1101/2022.08.09.503380

**Authors:** Zachary V. Johnson, Brianna E. Hegarty, George W. Gruenhagen, Tucker J. Lancaster, Patrick T. McGrath, Jeffrey T. Streelman

## Abstract

Social behaviors are essential for survival and reproduction and vary within and among species. We integrate single nucleus RNA-sequencing (snRNA-seq), comparative genomics, and automated behavior analysis to investigate a recently-evolved social “bower building” behavior in Lake Malawi cichlid fishes. We functionally profile telencephalic nuclei matched to 38 paired behaving/control individuals. Our data suggest bower behavior has evolved in part through divergence in a gene module selectively expressed in a subpopulation of glia lining the pallium. Downregulation of the module is associated with glial departure from quiescence and rebalancing of neuronal subpopulation proportions in the putative homologue of the hippocampus. We show further evidence that behavior-associated excitation of neuronal populations that project to the putative hippocampus mediate glial function and rebalancing. Our work suggests that bower behavior has evolved through changes in glia and region-specific neurogenesis, and more broadly shows how snRNA-seq can generate insight into uncharted behaviors and species.

## INTRODUCTION

Social behaviors vary tremendously within and among species, and they are disrupted in heritable human brain diseases (Johnson and Young 2017; Kennedy and Adolphs 2012). Many social behaviors are not expressed in standard laboratory models, and much progress in understanding the biological mechanisms of social behaviors has been made through work in diverse and non-traditional species systems (S. Juntti 2019; Gallant and O’Connell 2020; Laurent 2020; Keifer and Summers 2016; Brenowitz and Zakon 2015; Jourjine and Hoekstra 2021; Johnson and Young 2018). Different experimental traditions spanning genomics (C. R. Smith et al. 2008; Küpper et al. 2016; Lamichhaney et al. 2016; Bendesky et al. 2017; York et al. 2018; Pfenning et al. 2014; Dias and Walsh 2020; Stein et al. 2017), endocrinology (S. A. Juntti et al. 2016; Boender and Young 2020; Adkins-Regan 2013; O’Connell, Matthews, and Hofmann 2012; S. Ogawa et al. 2000; Heinrichs and Gaab 2007; Schiller, Meltzer-Brody, and Rubinow 2015), and circuit neuroscience (Gutzeit et al. 2020; Hung et al. 2017; Amadei et al. 2017; Anderson 2016; Gangopadhyay et al. 2021; Kohl et al. 2018; S. B. Nelson and Valakh 2015; Bachevalier and Loveland 2006) have contributed to our understanding of social behavior. However, we still have a poor understanding of the genetic and cellular pathways through which social behaviors vary and evolve. Discovering these gene-brain-behavior links is necessary to understand how neural circuit functions vary during social contexts.

Single cell omics technologies enable simultaneous profiling of many heterogeneous cell populations in any species with a reference genome, eroding important historical barriers that have faced investigation of new social behaviors and species systems. These technologies have already advanced our understanding of the brain (Tosches et al. 2018; Jerber et al. 2021; Raj et al. 2018; M. Zhang et al. 2021), however, to our knowledge only one study has used single cell omics to functionally profile the brain during behavior (Moffitt et al. 2018). Here we integrate single nucleus RNA-sequencing (snRNA-seq) with automated behavior analysis and comparative genomics to investigate the neurobiological substrates of a recently-evolved (<1 Mya) social bower construction behavior in Lake Malawi cichlid (*Cichlidae*) fishes. Cichlids are teleost (*Teleostei*) fishes, a group representing ∼40% of all living vertebrate species (Salzburger 2018). As teleosts, cichlids possess predicted homologues for ∼80% of human disease-associated genes (Howe et al. 2013). In the brain, teleosts and mammals share conserved neuronal and non-neuronal cell populations with conserved molecular, electrophysiological, morphological, transcriptional, and behavioral properties (O’Connell and Hofmann 2011b; Xie and Dorsky 2017; Elliott et al. 2017; Jurisch-Yaksi, Yaksi, and Kizil 2020). For example, the teleost telencephalon contains conserved cell populations that are thought to regulate social behaviors across diverse vertebrate lineages (O’Connell and Hofmann 2011b).

Lake Malawi is home to ∼800 cichlid species are behaviorally diverse (York et al. 2015; Baran and Streelman 2020; Ribbink et al. 1983; Johnson, Moore, et al. 2020; York et al. 2018) but genetically similar (Loh et al. 2008; Malinsky et al. 2018), thus representing a powerful system for investigating the neurogenetic basis of behavioral variation. In ∼200 species, males express bower construction behaviors during the breeding season, during which they repetitively spatially manipulate sand into species-specific structures for courtship and mating (York et al. 2015; Johnson, Arrojwala, et al. 2020; Long et al. 2020). Many species dig crater-like “pit” depressions while others build volcano-like “castle” elevations, and these behavioral differences are associated with genomic divergence in a ∼19 Mbp chromosomal region enriched for human disease-associated genes and genes that exhibit *cis*-regulated behavior-associated expression in the cichlid brain (York et al. 2018).

In this paper we investigate castle-building behavior in *Mchenga conophoros*, a Lake Malawi cichlid and an uncharted species in behavioral neuroscience. We use natural genetic differences among individuals to link single nuclei back to 38 paired behaving/control test subjects, enabling measurement of building-associated signals and simultaneous control for two additional biological variables that may influence brain gene expression: quivering, a courtship “dance” behavior, and relative gonadal mass. We first map the cellular diversity of the telencephalon and then investigate cell type-specific signatures of active castle-building behavior as well as genomic divergence associated with behavioral evolution. Our work shows how snRNA-seq profiling can generate converging lines of evidence for candidate genes, molecular signaling systems, cell populations, and brain regions underlying social behaviors in uncharted species systems.

## RESULTS

### Castle-building is associated with increased quivering behavior and gonadal physiology

We used an automated behavior analysis system (Johnson, Arrojwala, et al. 2020; Long et al. 2020) to monitor reproductive adult *Mchenga conophoros* males as they freely interacted with four reproductive adult females and sand (Fig. 1A). This system uses depth sensing to measure structural changes across the sand surface and action recognition to predict building and quivering (a stereotyped courtship “dance” behavior) from video data. We sampled pairs of males at the same time in which one male was actively castle-building within the past two hours (n=19) and the other was not (“control”, n=19; Fig. 1B-C). For each subject, we also recorded the gonadal somatic index (GSI), a measure of relative gonadal mass that is correlated with gonadal steroid hormone levels and social behaviors in cichlids (Maruska and Fernald 2010; Ramallo et al. 2015; Alward et al. 2019) (Table S1). The volume of sand displaced by males was positively correlated with the number of building events predicted from video data by action recognition (Fig. 1D). For simplicity, we combined depth and action recognition data into a single “Bower Activity Index” (BAI). Building males had greater BAIs, quivered more, and had greater GSIs (Fig. 1E-I) compared to controls. Taken together, these results are consistent with castle-building, like many social behaviors in nature, being embedded within a suite of behavioral and physiological changes tied to reproduction.

**Figure 1.**
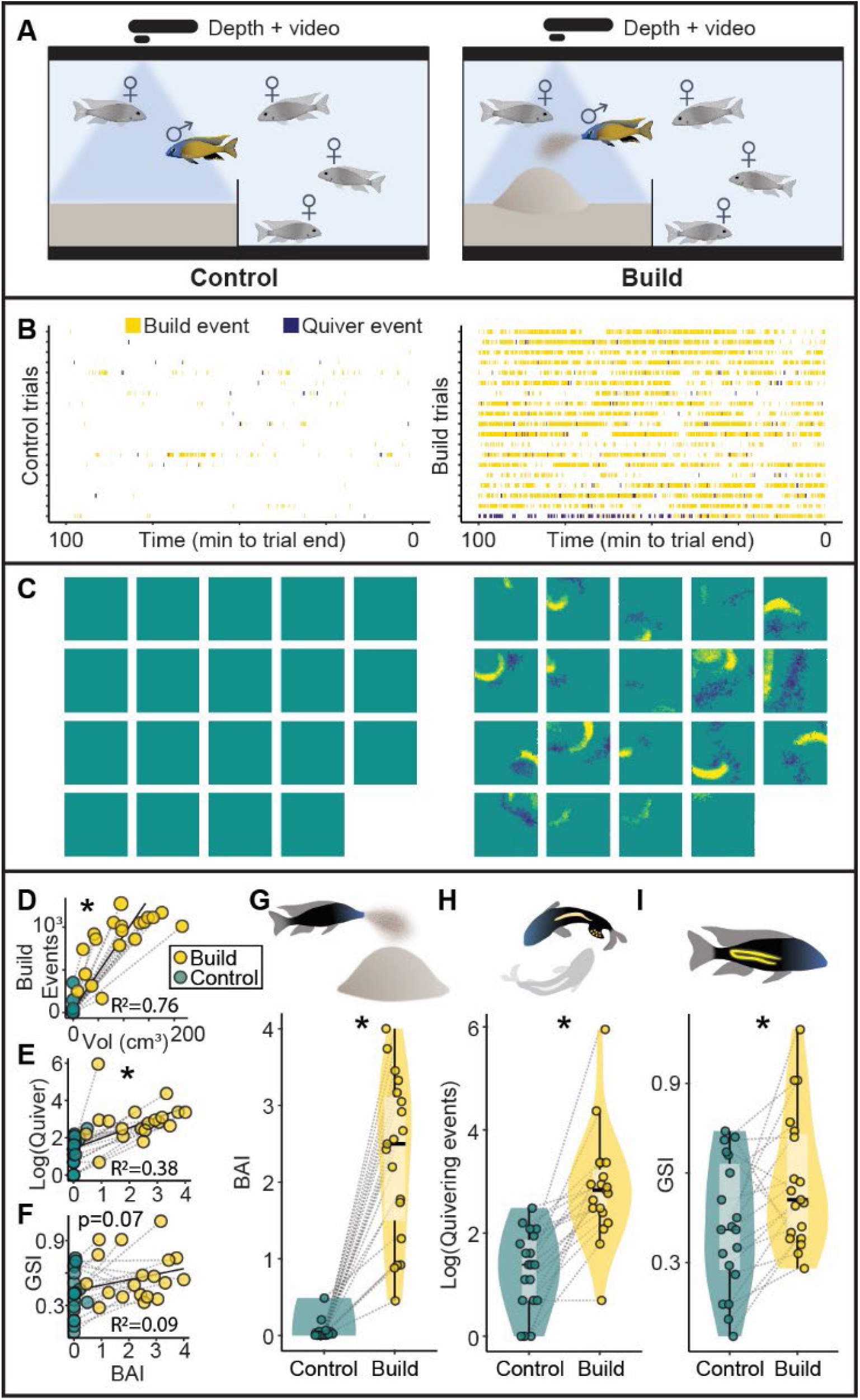
Castle-building is associated with increased quivering and relative gonadal mass. (A) Schematic of behavioral assay, 19 pairs of building (right) and control (left) males were sampled. Action recognition (B, yellow=building, blue=quivering, each trial is represented by a row, with pairs matched by row between left and right panels) and depth sensing (C, yellow=elevations, blue=depressions, each square represents total depth change for one trial, with pairs matched by row and column between left and right panels) revealed behavioral differences between building and control males. (D) Structural change measured through depth sensing (adjusted for body size) was strongly and positively correlated with building behaviors predicted through action recognition (p=8.15×10^−13^), and these measures were combined into a single Bower Activity Index (BAI, x-axis in E and F). BAI was positively correlated with quivering behaviors (E, p=3.35×10^−5^), and trended toward a positive correlation with GSI (F, p=0.07). Compared to controls, building males exhibited greater BAIs (G, 4.24×10^−8^), quivering (H, p=9.18×10^−6^), and GSIs (I, p=0.0142). Gray lines in panels D-I link paired building and control males.

### Telencephalic nuclei reflect major neuronal and non-neuronal cell classes

Telencephala (n=38) were combined into ten pools (n=5 behave, n=5 control, 3-4 telencephala/pool) for snRNA-seq (Fig. 2A). >3 billion RNA reads were sequenced and mapped to the Lake Malawi cichlid *Maylandia zebra* reference genome (Conte et al. 2019). 33,674 nuclei (∼900 nuclei/subject) passed quality control filters and were linked back to test subjects using genomic DNA. Coarse-grained clustering grouped nuclei into 15 “primary” (1°) clusters and finer-grained clustering grouped nuclei into 53 “secondary” (2°) clusters (ranging from 57-1,905 nuclei, Fig. 2B). Established marker genes revealed known neuronal and non-neuronal cell types (Fig. 2C), including excitatory (*slc17a6+*) and inhibitory (*gad2+*) neurons, oligodendrocytes and oligodendrocyte precursor cells (OPCs, *olig2+),* radial glial cells (RG, *fabp7+*), microglia, pericytes, and hematopoietic stem cells (Table S2). Unbiased analysis identified genes exhibiting nearly cluster-exclusive expression (Fig. 2D, top rows). Different clusters also exhibited preferential expression of genes encoding transcription factors (TFs; Fig. 2E-F) and neuromodulatory signaling molecules (Fig. 2G-H) that exhibit conserved neuroanatomical expression patterns in teleosts (Table S2). (Fig. 2I, Table S3). Cluster composition was relatively consistent across individuals. For clarity, we assigned each 1° cluster a numeric identifier (1-15) followed by a label indicating one or more of these cell classes (e.g. for radial glia, “_RG”). 2° cluster labels were rooted in these 1° labels, but with a second numeric identifier indicating the relative size within the corresponding “parent” 1° cluster (e.g. “4_GABA” is a 1° cluster expressing inhibitory neuronal markers, and “4.3_GABA” is the third largest 2° cluster within 4_GABA). Marker genes for every individual 1° and 2° clusters were independently enriched (q<0.05) for eight GO categories related to cell morphology, connectivity, conductance, and signal transduction (Table S4), supporting these as additional axes distinguishing clusters in this study. Cluster marker genes were also more strongly enriched for genes encoding conserved brain region-specific neurodevelopment/neuroanatomy-associated TFs (nTFs, n=43) and ligands (“ligands”, n=35) compared to neuromodulatory receptors (“receptors”, n=108, Table S5; receptors versus nTFs, p≤8.33×10^−4^ for both 1° and 2° clusters, FET; receptors versus ligands, p≤0.0068 for both; nTFs versus ligands, p≥0.75 for both, Fig. 2J), consistent with recent single cell RNA-seq (scRNA-seq) analyses of the mouse hypothalamus (Moffitt et al. 2018). Notably, several nTFs involved in dorsal-ventral patterning in early neural development exhibited striking polarity in expression across clusters (Fig. 2F). For example, *dlx* genes and *isl1* mark the ventral telencephalon while *emx* genes mark the dorsal telencephalon during the neurula stage (Sylvester et al. 2013), suggesting that transcriptional signatures of developmental patterning are present in adult neurons. Together these data may reflect organizing principles whereby transcriptional programs related to neurodevelopment and ligand synthesis are less labile, while neuromodulatory receptors are expressed more promiscuously across cell populations.

**Figure 2.**
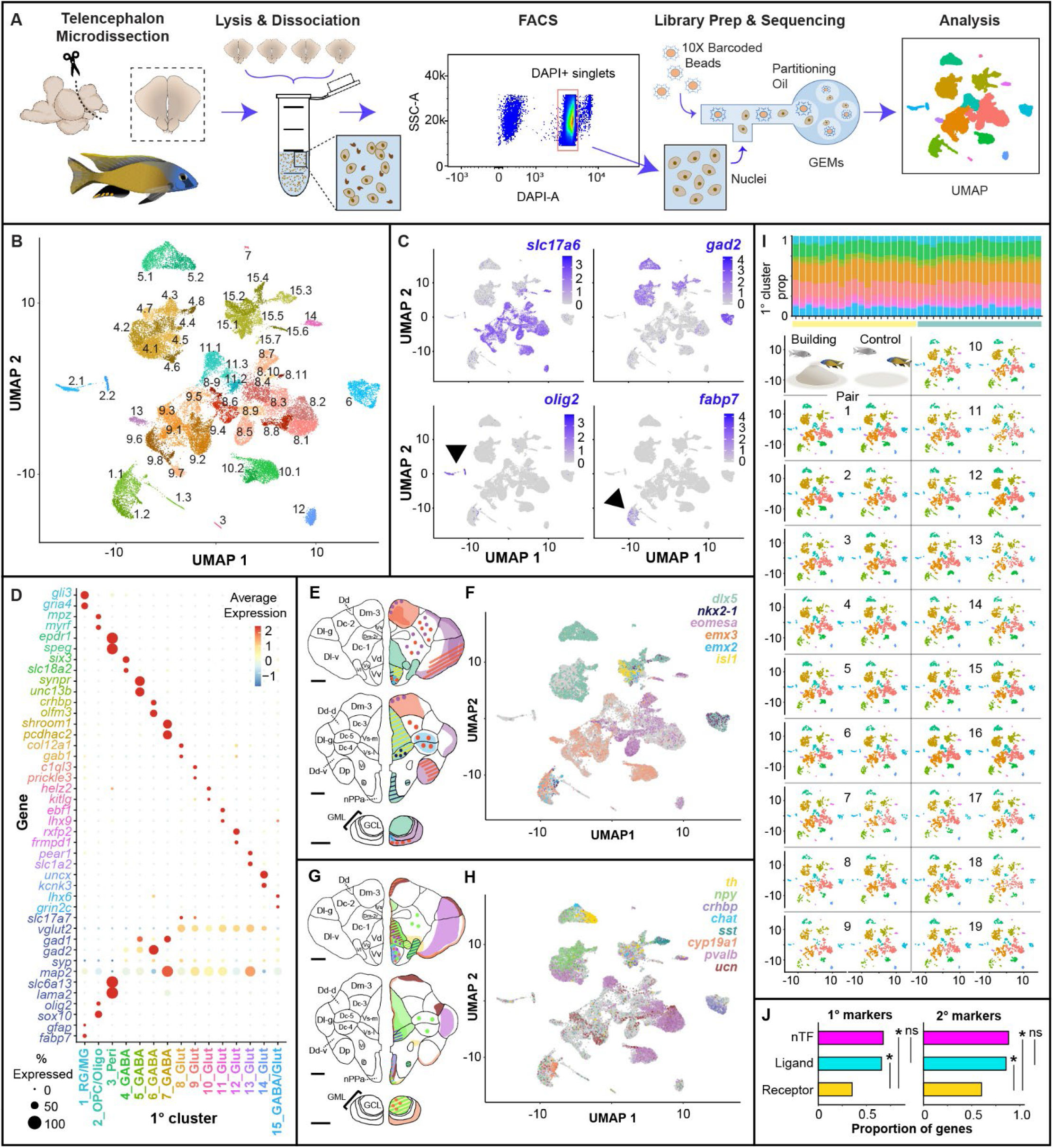
Molecular and cellular diversity of the cichlid telencephalon. (A) Schematic of experimental pipeline for snRNA-seq. (B) Nuclei cluster into 1° (n=15) and 2° (n=53) clusters. (C) Known marker genes reveal distinct clusters of excitatory neurons (*slc17a6*+), inhibitory neurons (*gad2*+), oligodendrocytes and oligodendrocyte precursor cells (*olig2*+), radial glia (*fabp7*+), as well as other less abundant cell types (not shown, see Table S2). (D) Clusters are distinguished by genes that exhibit near cluster-exclusive expression (top rows) as well as established cell type marker genes (bottom rows). Conserved nTFs (E, F) and ligands (and related genes; G, H) exhibit conserved neuroanatomical expression profiles in teleost fishes (E, G show schematic representations of conserved expression patterns), and show distinct expression in specific clusters. (I) Cluster proportions are consistent across 38 males (yellow and turquoise coded columns in top stacked bar chart represent building and control subjects, respectively). (J) nTF and ligand genes are differentially overrepresented among 1° and 2° cluster markers compared to receptor genes. Anatomical figures adapted with permission from Dr. Karen Maruska (Maruska et al. 2017).

### Building, quivering, and gonadal physiology are associated with signatures of neuronal excitation in distinct cell populations

To identify candidate cell populations that may regulate castle-building behavior, we first investigated transcriptional signatures of neuronal excitation. Neuronal excitation triggers intracellular molecular cascades that induce transcription of conserved immediate early genes (IEGs) (Lyons and West 2011), and mapping IEG expression is a strategy for identifying neuronal populations that are excited by specific stimuli or behavioral contexts (Guzowski et al. 2005). IEG transcripts tend to be recovered at lower levels compared to other genes in sc/snRNA-seq data (Y. E. Wu et al. 2017; Lacar et al. 2016; Moffitt et al. 2018). To better track these signals, we identified genes that were selectively co-transcribed with three established IEGs (*c-fos, egr1, npas4*) independently across 2° clusters. In total, we identified 25 “IEG-like” genes (Table S6), most (17/25, 68%) of which had previously been identified as IEGs, but eight of which have not (predicted homologues of human *DNAJB5, ADGRB1, GPR12, ITM2C, IRS2, RTN4RL2, RRAD*; Fig. 3A). These genes may include new markers of neuronal excitation.

**Figure 3.**
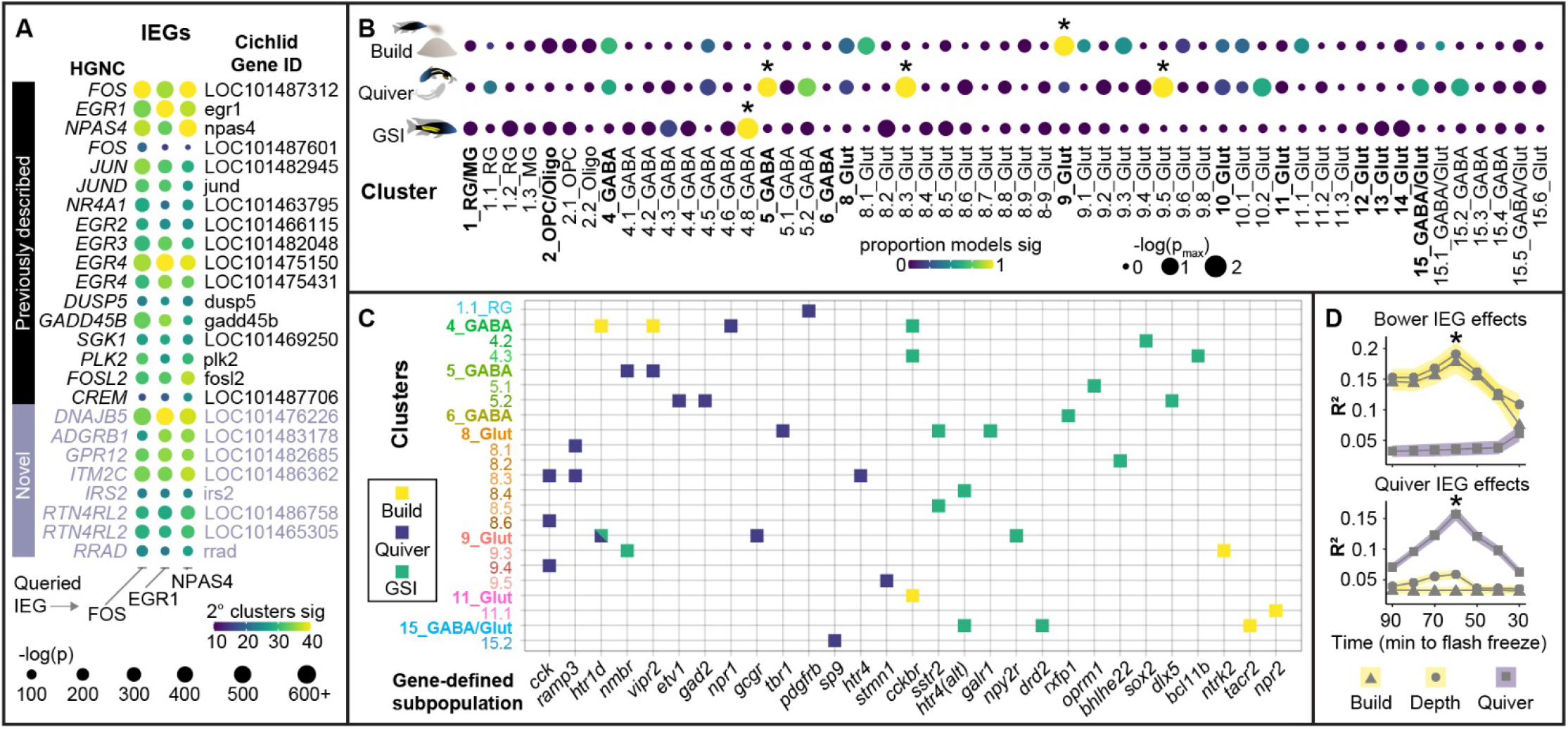
Distinct cell populations exhibit building-, quivering-, and gonadal-associated IEG expression. (A) 25 genes were selectively co-expressed with *c-fos, egr1*, and *npas4* across cell populations. (B) Building-, quivering-, and gonadal-associated IEG expression was observed in distinct clusters and (C) gene-defined populations (filled squares indicate significant effects, q<0.05). (D) IEG expression was most strongly associated with the amount of building (top) and quivering (bottom) behavior performed approximately 60 minutes prior to tissue freezing.

We assigned each nucleus an “IEG score,” equal to the number of unique IEG-like genes expressed. To disentangle building-, quivering-, and GSI-associated signals, we tested a sequence of models in which these variables competed in different combinations to explain variance in IEG score. Effects were considered significant if the raw p-value was significant (p<0.05) in every model and if the FDR-adjusted harmonic mean p-value (hmp_adj_) was significant across models (hmp_adj_<0.05) (Wilson 2019). Building was associated with increased IEG expression in 9_Glut (hmp_adj_=0.0016; Fig. 3B), a cluster with gene expression patterns reflective of Dd and Dc, two pallial brain regions (Martinelli et al. 2016). We also reasoned that some behaviorally-relevant populations may not align with clusters. For example, neuropeptides can diffuse to modulate distributed cell populations expressing their target receptors (Johnson and Young 2017), and other behaviorally relevant populations may represent a small proportion of one cluster. We therefore analyzed populations defined by nTF, ligand, and receptor genes, as well as a small set of additional genes of interest (n=17, “Other”, Table S5), both within clusters and regardless of cluster. IEG score was associated with building, quivering, and GSI in distinct cell populations (Fig. 3B; Table S6). Building was associated with IEG score in three populations defined regardless of cluster (*elavl4+, cckbr+, ntrk2*+), and in 4_GABA *htr1d+*, 4_GABA *vipr2+*, 15_GABA/Glut *tacr2*+, 11_Glut *cckbr*+, and 11.1_Glut *npr2*+ nuclei (Fig. 3C), consistent with a role for these molecular systems in the neural coordination of building. Quivering was associated with IEG score in 5.2_GABA *etv1+* nuclei, a subpopulation strongly expressing a suite of dopamine (e.g. *etv1, th, dat, vmat*) and progenitor (e.g. *etv1, pax6*) neuron marker genes that are known to be expressed in the olfactory bulb granule cell layer, a region in which new dopaminergic neurons are born in adult teleosts. These data are consistent with previous work showing activation of olfactory and dopaminergic circuitry during courtship in diverse systems (Keleman et al. 2012; van Furth, Wolterink, and van Ree 1995; Ishii and Touhara 2019; Louilot et al. 1991; Johnson and Young 2015). Building- and quivering-associated IEG signals were most strongly associated with behavior expressed approximately 60 minutes prior to sample collection, consistent with previously reported IEG nuclear RNA time courses (Lacar et al. 2016) and further reinforcing their behavioral significance (Fig. 3D).

### A minority of neuronal populations account for the majority of building-associated gene expression

Social behaviors have been linked to large changes in brain gene expression in diverse lineages (Robinson, Fernald, and Clayton 2008; Baran and Streelman 2020; Patil et al. 2021; York et al. 2018), but the underlying cell populations driving these effects are not well understood. We performed an unsupervised analysis to identify differentially expressed genes (DEGs) in specific clusters. A relatively small subset of neuronal clusters accounted for a disproportionate number of building-associated DEGs (bDEGs), a pattern that was also true of quivering-associated DEGs (qDEGs) and gonadal-associated DEGs (gDEGs; Fig. 4A; Table S7). bDEGs were overrepresented in three excitatory neuronal clusters (8_Glut, 9_Glut, 10_Glut; q≤1.83×10^−4^ for all), qDEGs were overrepresented in two neuronal clusters (15_GABA/Glut, 11_Glut, q≤0.036 for both), and gDEGs were overrepresented in one inhibitory neuronal cluster (5_GABA, q=1.30×10^−5^). bDEGs were overrepresented in a suite of aligned 2° clusters (q≤6.69×10^−4^ for all), qDEGs were overrepresented in 15.2_GABA, 8.1_Glut, and 8.6_Glut (q≤0.0074 for all), and gDEGs were overrepresented in 8.3_Glut and 8.4_Glut (q≤0.039 for both). Thus, distinct clusters were overrepresented for bDEGs, qDEGs, and gDEGs. Interestingly, despite these non-overlapping signals across clusters, a substantial set of bDEGs, gDEGs, and qDEGs were the same individual genes (n=81), consistent with behavior and gonadal hormones recruiting common transcriptional programs in distinct populations (Fig. 4B). These results highlight a small set of 1° and 2° neuronal clusters as candidate regulators of castle-building behavior.

**Figure 4.**
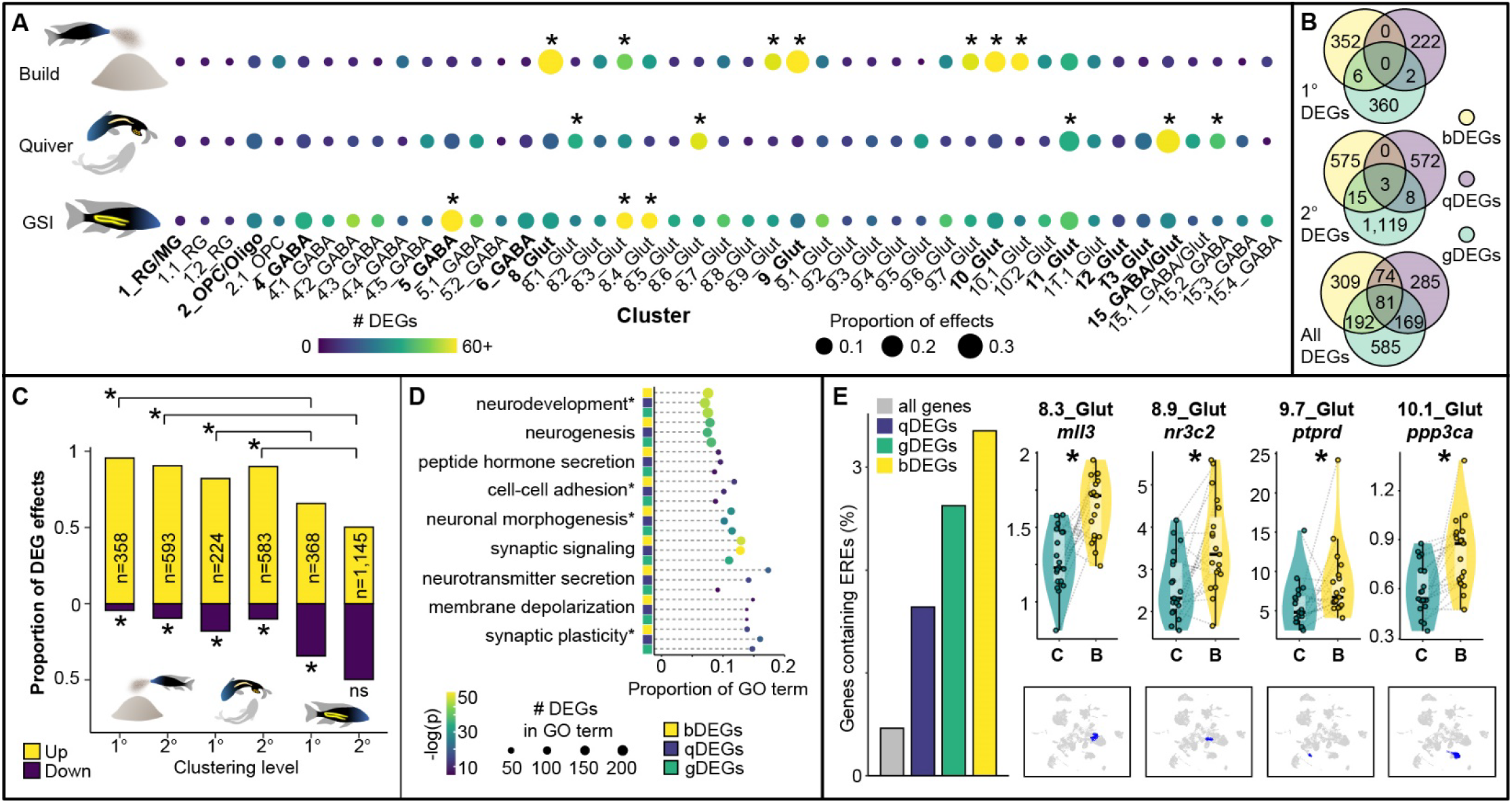
Building, quivering, and GSI are associated with distinct patterns of cell type-specific gene expression. (A) Distinct 1° and 2° clusters show a disproportionate number of bDEGs, qDEGs, and gDEGs. (B) A set of 81 genes exhibits building-, quivering, and gonadal-associated expression in largely non-overlapping clusters. (C) Behavior-associated gene expression is driven by upregulation, whereas gonadal-associated gene expression is driven by a balance of up- and downregulation. (D) bDEGs, qDEGs, and gDEGs are enriched for GO terms related to synaptic structure, function, and plasticity; neurotransmission; and neurogenesis. (E) bDEGs, qDEGs, and gDEGs are enriched for EREs. Violin plots show cluster-specific ERE-containing bDEG effects and feature plots below show the clusters (blue) in which each effect was observed. GO terms followed by asterisks are abbreviated.

Behavior-associated DEGs exhibited a strong direction bias, and were predominantly upregulated in both 1° and 2° clusters (p≤1.39×10^−12^ for all, Fig. 4C). In contrast, gDEGs tended more modestly toward upregulation in 1° clusters (1° gDEG effects, p=2.09×10^−5^) and were not directionally biased in 2° clusters (2° gDEG effects, p=0.92). Upregulated bDEGs, qDEGs, and gDEGs were each independently enriched for a large number of the same GO terms (q<0.05 for 499 GO Biological Processes, 147 GO Cellular Components, and 111 GO Molecular Functions), the strongest of which were related to synaptic transmission and plasticity (e.g. “synaptic signaling,” q≤3.54×10^−50^ for all; “regulation of synaptic plasticity,” q≤1.83×10^−18^ for all) or cell differentiation and neurogenesis (e.g. “nervous system development,” q≤6.93×10^−47^ for all; “neurogenesis,” q≤4.49×10^−35^ for all; “cell morphogenesis involved in neuron differentiation,” q<6.96×10^−29^ for all; Fig. 4D), suggesting behavior- and gonadal-associated regulation of synaptic function and cell morphogenesis.

### Estrogen response elements are enriched in behavior- and gonadal-associated differentially expressed genes

Estrogen regulates social behavior in diverse species and has been linked to both neuronal excitability and neurogenesis (Diotel et al. 2013; Duarte-Guterman et al. 2015; Kelly and Rønnekleiv 2009; Sarkar et al. 2008). Estrogen can also regulate gene expression by binding to estrogen receptors (ERs), forming a complex that translocates into the nucleus and acts as a TF by binding to Estrogen Response Elements (EREs) in DNA (Klinge 2001; Amenyogbe et al. 2020). bDEGs, gDEGs, and qDEGs were independently enriched for EREs, consistent with a role for estrogen in modulating behavior- and gonadal-associated gene expression (p≤2.92×10^−4^ for all; Fig. 4E; ERE-containing gene list in Table S8). ERE-containing bDEGs (n=22 unique genes) were most strongly enriched for GO terms including “modulation of chemical synaptic transmission” (top GO Biological Process, q=2.30×10^−4^) and “Schaffer collateral - CA1 synapse” (top Cellular Component, q=2.22×10^−5^), consistent with building-associated estrogenic regulation of synaptic function. These data support a role for estrogen in castle-building behavior.

### Castle-building is associated with neuronal rebalancing in the putative fish hippocampus

The enrichment of neurogenesis-related GO terms among bDEGs motivated us to further investigate building-associated neurogenesis. During neurogenesis, new neurons differentiate into specific neuronal populations (Mira and Morante 2020; Götz and Huttner 2005), and we therefore reasoned that building-associated neurogenesis may result in build-associated changes in the relative proportions of specific neuronal populations. Analysis of cluster-specific proportions revealed building-associated increases in the relative proportion of 8.4_Glut (q=0.013; Fig. 5A,B) and decreases in the relative proportion of 8.1_Glut (q=7.67×10^−4^; Fig. 5A,C). The relative proportions of 8.4_Glut and 8.1_Glut were negatively correlated across subjects, such that greater proportions of 8.4_Glut predicted lesser proportions of 8.1_Glut (R=-0.50, p=0.0012; Fig. 5D). Notably, 8_Glut was distinguished by markers of the lateral region of the dorsal telencephalon (Dl; Table S2), a brain region that is important for spatial learning, memory, and behavior in other fish species. Dl is the putative fish homologue of the mammalian hippocampus, a region in which adult neurogenesis regulates spatial learning and memory (Clark et al. 2008; Clelland et al. 2009).

**Figure 5.**
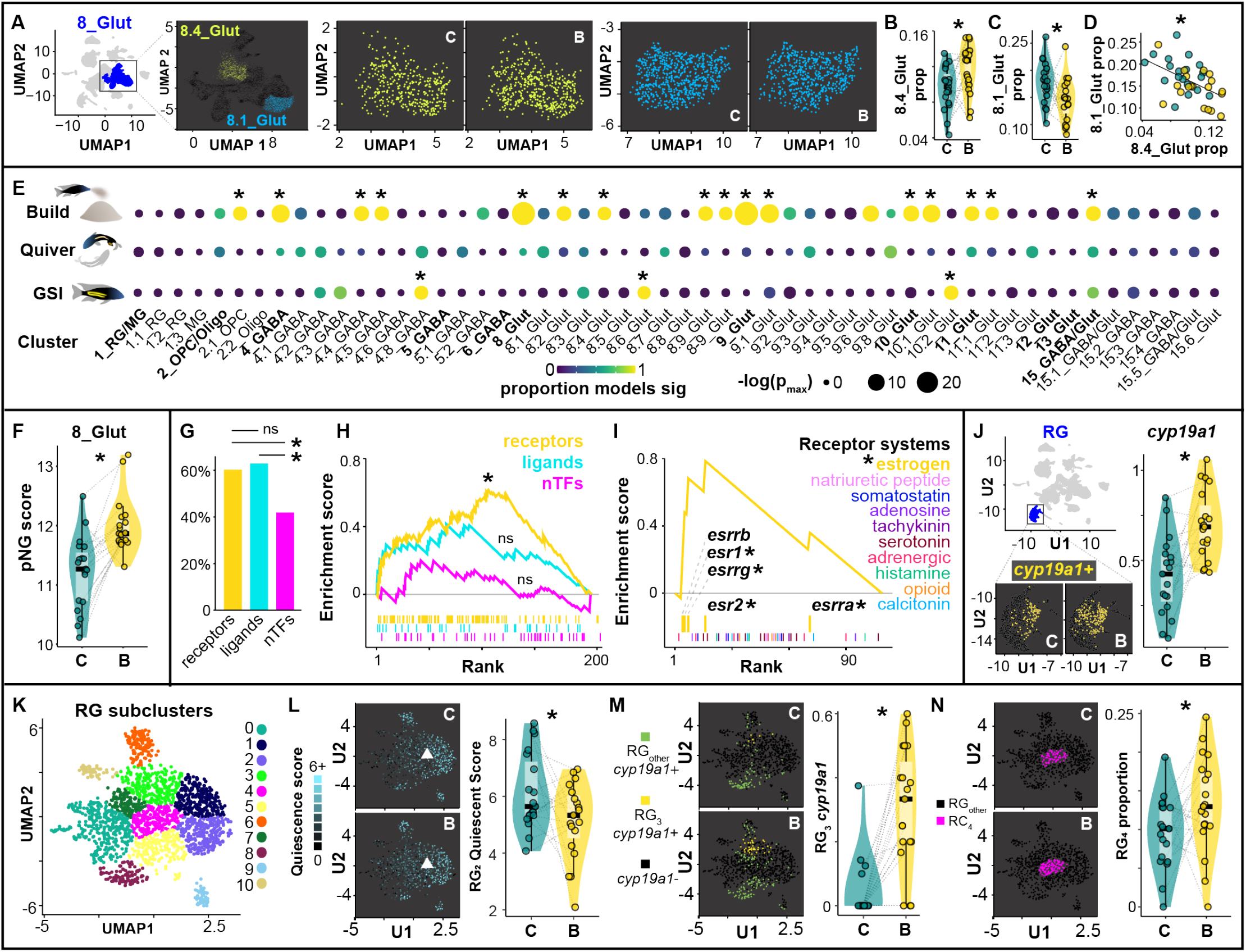
Behavior is associated with signatures of neurogenesis in neurons and glia. (A-C) Building is associated with a shift in the relative proportions in 8.4_Glut and 8.1_Glut, and (D) the relative proportions of these two clusters is strongly correlated across individuals. (E) Building, but not quivering, is associated with increased pNG expression in a large set of 1° and 2° clusters, whereas GSI is associated with increased and decreased pNG expression in just three 2° clusters. (F) The most significant building-associated pNG expression is observed in 8_Glut. (G) Gene-defined populations that exhibit building-associated pNG expression are disproportionately defined by genes encoding receptors and ligands. (H) The strongest building-associated pNG expression tends to occur in populations defined by neuromodulatory receptors, (I) particularly in ER-expressing populations. (J) RG exhibit building-associated *cyp19a1* expression. (K) Reclustered RG subpopulations show building-associated (L) signatures of decreased quiescence (RG_2_), (M) *cyp19a1* expression (RG_3_), and (N) increases in proportion (RG_4_).

### Castle-building is associated with increased expression of genes that positively regulate neurogenesis

To further investigate building-associated neurogenesis, we identified 87 genes with the GO annotation “positive regulation of neurogenesis” in both zebrafish and mice (“proneurogenic” genes, pNGs, Table S9) and analyzed their expression across clusters and gene-defined populations. Building was associated with increased pNG expression in six 1° clusters (8_Glut, 9_Glut, 10_Glut, 11_Glut, 15_GABA/Glut, 4_GABA) and ten aligned 2° clusters (including 8.4_Glut; hmp_adj_≤0.020 for all; Fig. 5E). The most significant building-associated pNG expression was observed in 8_Glut (Fig. 5F, hmp_adj_=1.23×10^−17^), and pNG expression in 8.4_Glut specifically was positively associated with its relative proportion (R=0.33, p=0.041). In contrast to building, gonadal-associated pNG expression was increased in 10.2_Glut (hmp_adj_=0.010) and decreased in 4.8_GABA (hmp_adj_=0.0048), and quivering was not associated with pNG expression in any 1° or 2° clusters (Fig. 5E). Notably, the magnitude of effect (β) estimates for building-associated pNG expression in 2° clusters were always greater than in their “parent” 1° clusters, and many gene-defined subpopulations within clusters exhibited stronger building-associated pNG expression than their parent 1° or 2° clusters. For example, within 15_GABA/Glut, building-associated pNG estimates were >3x greater in subpopulations defined by expression of *adra2b* (β_*cond=*_=0.188) and *esr2* (β_*cond*_=0.154) compared to 15_GABA/Glut as a whole (β_*cond*_=0.048). Among 2° clusters, the most extreme cases of this pattern included 8.2_Glut *drd4+*, 8.4_Glut *htr4+*, 9.1_Glut *sstr5+*, 9.6_Glut *htr4+*, 10.1_Glut *ntrk2+* nuclei, and 11.1_Glut *ntrk2+* nuclei (hmp_adj≤_0.018 for all). Among populations defined regardless of cluster, those exhibiting building-associated pNG expression were disproportionately defined by neuromodulatory receptor and ligand genes versus nTFs (receptors versus nTFs, q=0.011; ligands versus nTFs, q=0.017; FET; Fig. 5G), and those exhibiting the strongest building-associated pNG expression (β) effects were disproportionately defined by neuromodulatory receptor genes (q=0.011; Fig. 5H), and by ERs in particular (q=0.034; Fig. 5I), consistent with a large body of literature supporting relationships between estrogen and neurogenesis (Diotel et al. 2013; Duarte-Guterman et al. 2015). These results highlight specific molecular signaling systems (e.g. estrogen, serotonin, TrkB) that may be involved in building-associated neurogenic changes.

### Building is associated with changes in glial cell biology

Radial glia (RG) are the primary source of new neurons in adult teleosts (Ganz and Brand 2016), and we therefore reasoned that signatures of neurogenesis may be downstream effects of changes in RG function. We first investigated building-associated gene expression within radial glia (1.1_RG and 1.2_RG pooled). We identified 25 bDEGs that were collectively enriched for “neuron development” (top GO Biological Process, q=8.18×10^−4^) as well as “astrocytic glutamate-glutamine uptake and metabolism” (top Pathway, q=0.0010) and “synapse” (top GO Cellular Component, q=0.0015). RG bDEGs included *cyp19a1* (upregulated; Fig. 5J), the gene encoding aromatase, an enzyme that converts testosterone to brain-derived estrogen and has been previously linked to RG function and neurogenesis (Pellegrini et al. 2016).

RG can occupy distinct functional states including quiescence, cycling, and neuronal differentiation (Jurisch-Yaksi, Yaksi, and Kizil 2020; Adolf et al. 2006; Labusch et al. 2020). We re-clustered RG (independently of non-RG nuclei) into 11 subclusters (RG_0_-RG_10_; Fig. 5K) and assigned each nucleus a quiescence, cycling, and neuronal differentiation score based on established marker genes (Table S10), and analyzed building-associated differences in these scores across subclusters. Building was associated with decreased quiescence score in RG_2_ (hmp_adj_=0.010; Fig. 5L), but was not associated with quiescent, cycling, or neuronal differentiation score in any other subcluster. Analysis of building-associated gene expression across subclusters further revealed that 19/61 subcluster bDEGs were in RG_2_, and 18/19 effects reflected building-associated downregulation. The strongest enrichment hit for RG_2_ bDEGs was GO Cellular Component “postsynaptic Golgi apparatus” (q=0.0011). *cyp19a1* was excluded from analysis in several subclusters because it was not detected in all build-control pairs; however, a targeted analysis revealed that building-associated increases in *cyp19a1* were driven by RG_3_ (hmp_adj_=0.018; Fig. 5M), a subpopulation distinguished by *lhx5* and *gli3*, both nTFs that regulate neurogenesis in mammals (Y. Zhao et al. 1999; Hasenpusch-Theil et al. 2018). Lastly, because RG subclusters strongly aligned with functional states, we reasoned that building-associated transitions in RG function may also manifest as building-associated changes in subcluster proportions. Indeed, building was associated with an increase in the relative proportion of RG_4_ (q=0.0017; Fig. 5N), a subcluster positioned in UMAP space between nuclei expressing markers of quiescence and nuclei expressing markers of cycling. These data support building-associated changes in radial glial cell biology, and highlight RG_2_, RG_3_, and RG_4_ as candidate RG subpopulations involved in building-associated and RG-mediated neurogenesis.

### Genes that have diverged in castle-building lineages are upregulated in reproductive contexts

Castle-building behavior has previously been linked to a ∼19 Mbp region on Linkage Group 11 (LG11), within which genetic variants have diverged between closely-related castle-building and pit-digging lineages (York et al. 2018; Patil et al. 2021). Our follow up comparative genomics analyses identified 165/756 genes in this region that also showed signatures of divergence between castle-building lineages and more distantly-related “mbuna” species that do not build bowers (“castle-divergent” genes, CDGs; Fig. 6A; Table S11). Thus, CDGs represent a subset of genes bearing strong genomic signatures of castle-building evolution across Lake Malawi species. CDGs were expressed at higher levels in the telencephalon compared to neighboring genes in the same 19Mbp region (∼2.9x greater expression, permutation test, p=1.42×10^−5^) and compared to other genes throughout the genome (∼2.6x greater expression, p=1.77×10^−6^). CDGs were also overrepresented among 1° and 2° cluster markers (versus neighboring LG11 genes, p≤1.66×10^−9^ for both; versus all other genes, p≤1.43×10^−11^ for both, FET), and among upregulated bDEGs, qDEGs, and gDEGs (versus neighboring LG11 genes, p≤0.0044 for all; versus all other genes, p≤0.0066 for all, FET; Fig. 6B). These data support the behavioral significance of CDGs in the telencephalon, and suggest that castle-building evolution has targeted genes that are selectively upregulated during reproductive contexts.

**Figure 6.**
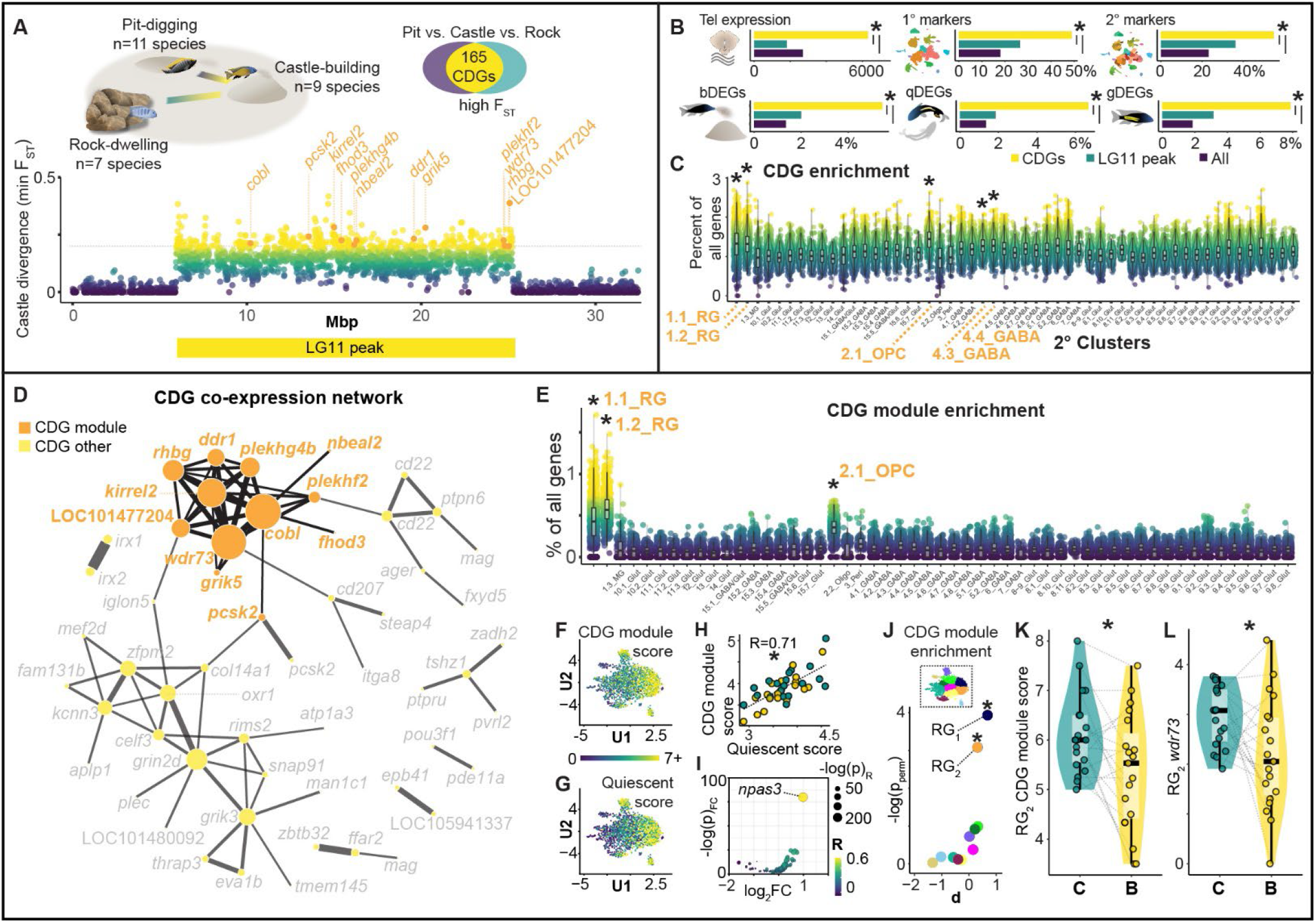
Genomic signatures of castle-building evolution link behavior, radial glial function, and hippocampal-like neuronal rebalancing. (A) Comparative genomics identifies 165 CDGs (CDG module genes labeled in orange). (B) CDGs are enriched in the telencephalon, among 1° and 2° cluster marker genes, and among bDEGs, qDEGs, and gDEGs. (C) CDGs are most strongly enriched in non-neuronal populations (y-axis shows the percentage of genes expressed that were CDGs). (D) A “CDG module” (orange) contains 12 CDGs that are strongly co-expressed across nuclei. (E) The CDG module is most strongly enriched in radial glia. (F,G) CDG module expression across radial glial subclusters mirrors expression of quiescent markers. (H) Expression of the CDG module is positively correlated with expression of radial glial quiescence markers. (I) *npas3* shows strong, positive, outlier co-expression with the CDG module. (J) RG_1_ and RG_2_ are enriched for the CDG module. (K) RG_2_ exhibits building-associated decreases in expression of the CDG module and *wdr73* in particular (L).

### Castle-divergent genes are enriched in quiescent radial glial subpopulations

CDGs were most strongly enriched in non-neuronal (2.1_OPC, 1.1_RG, and 1.2_RG), followed by neuronal (4.3_GABA and 4.4_GABA) clusters and gene-defined populations (5.2_GABA *th+*, and 9_Glut *hrh3+*; Fig. 6C; Table S12). We hypothesized that co-upregulation of subsets of CDGs in the same nuclei may drive cluster-specific enrichment patterns. A WGCNA (Langfelder and Horvath 2008) based analysis revealed a module of 12 CDGs that were more strongly co-expressed than other CDGs (stronger correlation coefficients, Welch t-test, p=8.83×10^−14^; stronger silhouette widths, Welch t-test, p=0.016; Fig. 6D). Across clusters, this module was most strongly enriched in 1.2_RG (p_perm_=0, Cohen’s d=4.22), and was less strongly enriched in 1.1_RG (Cohen’s d=2.86; Fig. 6E), suggesting differences in expression among RG subpopulations (Table S13). Within RG, CDG module expression was positively associated with quiescent score (Fig. 6F-H; R=0.34, p=3.21×10^−52^; p_perm_=0); and was negatively associated with cycling score (R=-0.089, p=9.90×10^−5^; p_perm_=0) and neuronal differentiation score (R=-0.065, p=0.0048; p_perm_=0). Analysis of co-expression between the module and known TFs (n=999) identified *npas3* as an outlier that was most strongly co-expressed TF with the CDG module (Fig. 6I; Table S14; R=0.47, q=3.19×10^−100^). *npas3* suppresses proliferation in human glioma, is strongly expressed in quiescent neural stem cells, and is downregulated during hippocampal neurogenesis in mice (Moreira et al. 2011; Shin et al. 2015). Among RG subclusters, the module was selectively enriched in RG_1_ (p_perm_=0.0196) and RG_2_ (p_perm_=0.046; Fig. 6J; Table S13), both of which selectively expressed genetic markers of RG quiescence. Together these data support that CDG module expression is positively related to RG quiescence.

### A subpopulation of glia links genome evolution to hippocampal-like neuronal rebalancing

Building was associated with a decrease in CDG module score in RG_2_ (hmp_adj_=0.027; Fig. 6K), and an increase in CDG module score in RG_8_ (hmp_adj_=0.010). The only individual CDG module gene for which we detected building-associated expression was *wdr73*, which showed building-associated downregulation in RG_1_ and RG_2_ (hmp_adj_≤4.54×10^−89^ for both; RG_2_ effect in Fig. 6L). These data raise the possibility of a building-associated downregulation of the CDG module and an exit from quiescence in RG_2_. We hypothesized that a building-associated exit from quiescence in RG_2_ may contribute to building-associated neuronal rebalancing between 8.4_Glut and 8.1_Glut. Consistent with this, the 8.4_Glut:8.1_Glut ratio was predicted by RG_2_ CDG module score (R=-0.52, p=6.91×10^−4^), *wdr73* expression (R=-0.62, p=3.31×10^−5^), quiescent score (R=-0.42, p=0.0094), and *npas3* expression (R=-0.52, p=8.20×10^−4^). All of these relationships were evident within building males only (8.4_Glut:8.1_Glut ratio versus RG_2_ CDG module score, R=-0.51, p=0.024; quiescent score, R=-0.42, p=0.059; versus RG_2_ *wdr73* expression, R=-0.59, p=0.0074; *npas3* expression R=-0.65, p=0.0027) but not within controls (p≥0.14 for all). In contrast, none of these relationships were evident in RG_1_, regardless of whether the analysis was conducted across all subjects (p≥0.13 for all) or restricted to building males (p≥0.074 for all). Together these data are consistent with a role for RG_2_ in neuronal rebalancing.

In teleost fishes, anatomically distinct RG subpopulations vary in function and supply new neurons to distinct brain regions (Fig. 7A). We hypothesized that if RG_2_ was involved in 8.4_Glut:8.1_Glut neuronal rebalancing, then its anatomical distribution should be consistent with supplying new neurons to brain regions within which 8.4_Glut nuclei reside. Spatial profiling revealed that 8.4_Glut and 8.1_Glut respectively mapped to ventral and dorsal Dl-v, a pallial subregion within Dl, the putatitve hippocampal homologue in fish (Fig. 7B-E). Thus, 8.4_Glut and 8.1_Glut both mapped to dorsolateral pallial regions that receive new neurons from RG lining the pallial ventricular zone. RG_2_ was anatomically positioned along the pallial but not subpallial ventricular zone (Fig. 7B-E), consistent with a potential to supply new neurons to Dl and other pallial regions. Together these data were consistent with a relationship between building-associated expression of the CDG module in RG_2_ and neuronal rebalancing in 8.4_Glut and 8.1_Glut.

**Figure 7.**
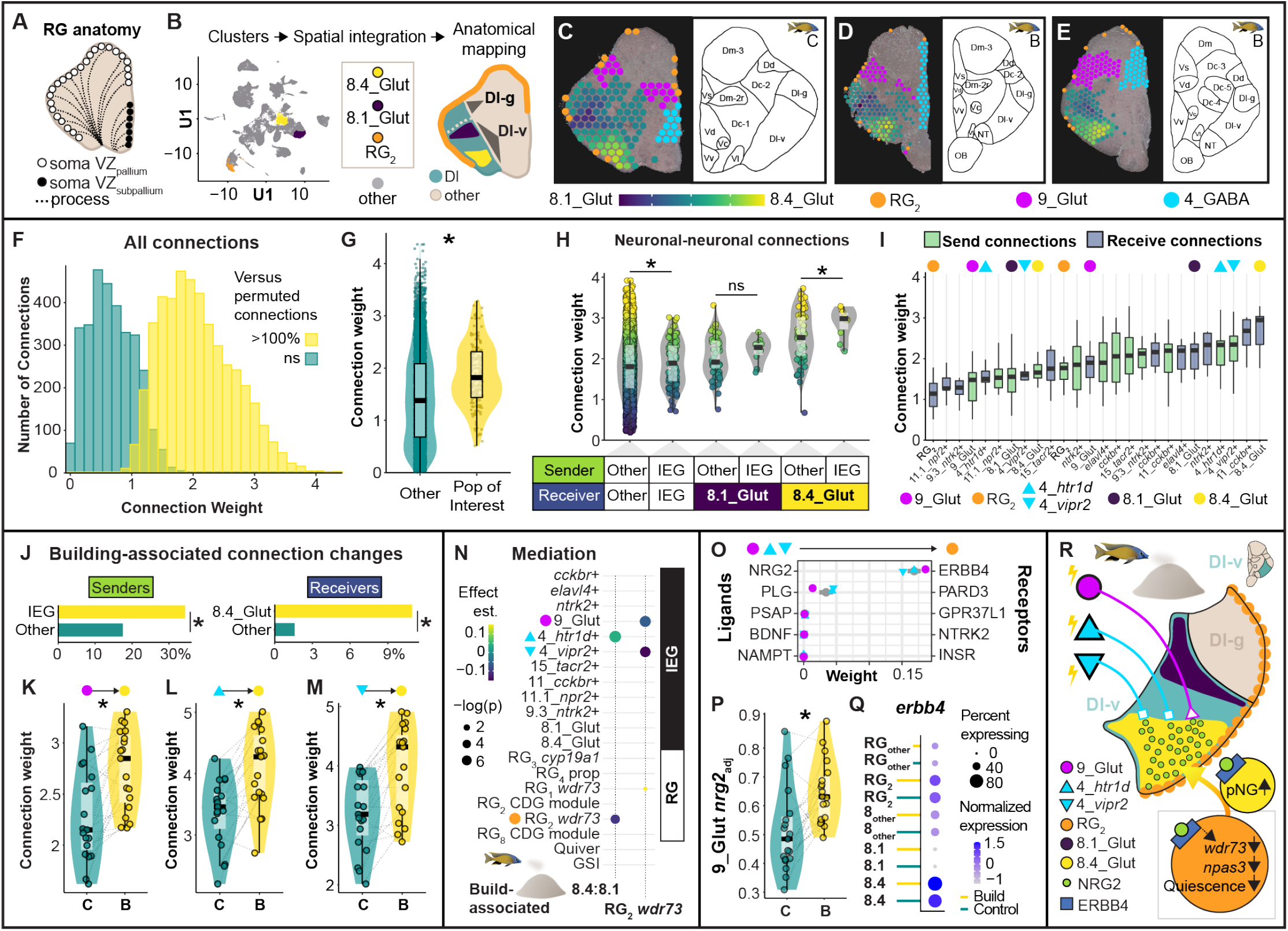
A circuit model for behavior-associated cellular reorganization in hippocampal-like Dl-v. (A) RG differ in morphology, function, and anatomical distribution (e.g. pallial versus subpallial ventricular zones). (B) Spatial profiling enables neuroanatomical mapping of RG_2_, 8.1_Glut, 8.4_Glut, and additional neuronal populations of interest, as illustrated in three individuals (C-E). RG_2_ (orange) aligns with the pallial but not subpallial ventricular zone, and 8.1_Glut versus 8.4_Glut aligns with dorsal versus ventral Dl-v, respectively. (F) Cell-cell communication analysis of randomly permuted populations separates bi-modally distributed connections among real populations. (G) Connection weights among populations of interest are stronger than among other populations. (H) Connection weights among build-IEG+ populations are greater than among other neuronal populations, and connection weights and between build-IEG+ populations (senders) and 8.4_Glut (receiver) are greater compared to other neuronal populations (senders) and 8.4_Glut (receiver). (I) Connections to 8.4_Glut (receiver) were greater than any other type of connection among populations of interest. (J) Connections exhibiting building-associated increases in strength were enriched for build-IEG+ senders and for 8.4_Glut as a receiver. (K-M) Connections from 9_Glut, 4_GABA *htr1d*+, and 4_GABA *vipr2*+ to 8.4_Glut all exhibit building-associated increases in weight. (N) Regularized multiple mediation analysis supports RG_2_ *wdr73* expression and 4_GABA *htr1d*+ IEG expression as mediators of 8.4_Glut:8.1_Glut neuronal rebalancing, as well as 9_Glut and 4_GABA *vipr2*+ IEG expression as mediators of RG_2_ *wdr73* expression. (O) NRG2-ERBB4 is the strongest cell-cell molecular signaling pathway identified between 9_Glut, 4_GABA *htr1d*+, and 4_GABA *vipr2*+ (senders) and RG_2_ (receiver). (P) *nrg2* shows building-associated upregulation in 9_Glut. (Q) *erbb4* shows preferential expression in both RG_2_ and 8.4_Glut. (R) A circuit model for how neural activity and RG_2_ *wdr73* expression may coordinate building-associated neuronal rebalancing in Dl-v.

### Populations excited during building may project to the putative fish hippocampus

In the mammalian hippocampus, the activity of local circuits and incoming projections regulate differentiation of glial cells into new neurons (Pardal and López Barneo 2016; Song et al. 2016). We reasoned that neural activity may similarly regulate building-associated mobilization of RG_2_ and neuronal rebalancing. We used CellChat (Jin et al. 2021) to investigate possible connections among 1° clusters, 2° clusters, RG subclusters, and nine gene-defined populations that, in addition to 9_Glut, showed signatures of building-associated excitation (“build-IEG+”; Table S15). Briefly, this tool estimates the molecular potential for connection (“weight”) between cell populations using known cell-cell adhesion and ligand-receptor binding proteins. Unlike most other tools, CellChat increases robustness by additionally accounting for heteromeric complexes and interaction mediator proteins (Dimitrov et al. 2022). As added control, we compared connections between pairs of populations to connections between randomly permuted cell populations of the same size, enabling us to identify connection weights that were greater than 100% of permuted results (Fig. 7F). Weights among neuronal populations of interest (8.1_Glut, 8.4_Glut, and build-IEG+ populations) were greater than weights among other neuronal populations (Fig. 7G; p=0.03, Welch two-sample t-test) and among permuted populations (all >100% of permuted connections). Neuronal populations of interest differed in “sending” (p=0.0016, Kruskal-Wallis rank sum test) and “receiving” (1.48×10^−14^) weights, and 8.4_Glut (receiver) had the greatest weights of any sender or receiver (Fig. 7H-I). Compared to other neuronal populations, build-IEG+ populations had greater sending weights to 8.4_Glut (p=0.027, Welch two-sample t-test), but not to 8.1_Glut (p=0.25; Fig. 7H). These data support a model whereby neuronal populations that fire during building project preferentially to 8.4_Glut neurons.

Neuronal firing can increase the strength of synaptic connections. ∼2% (64/3,136) of neuronal-neuronal connections exhibited building-associated changes in weight, with building males exhibiting stronger weights in every case. These connections were enriched for build-IEG+ senders (22/64, p=0.0014, FET) and 8.4_Glut as a receiver (7/64, p=1.11×10^−4^), but not for build-IEG+ receivers, 8.4_Glut as a sender, or 8.1_Glut as a sender or receiver (p≥0.50 for all; Fig. 7J). Build-IEG+ sender connections that showed building-associated change were enriched for 8.4_Glut as a receiver (4/22, p=3.24×10^−4^), and 8.4_Glut receiver connections that showed building-associated changes were enriched for build-IEG+ populations as senders (4/7, p=0.015). These patterns were driven by senders 9_Glut (Fig. 7K; hmp_adj_=0.0028), 4_GABA *htr1d+* (Fig. 7L; hmp_adj_=0.0081), 4_GABA *vipr2*+ (Fig. 7M; hmp_adj_=0.0027), *ntrk2+* (hmp_adj_=0.011) and 8.4_Glut (receiver). These data highlight specific populations that may project to Dl-v and fire during building.

### A behavioral circuit model for activity- and glial-dependent neurogenesis in the putative hippocampus

To investigate relationships among building-associated neural activity, changes in RG biology, and 8.4_Glut:8.1_Glut neuronal rebalancing, we used a regularized (LASSO) multiple mediation approach. Briefly, this tested if the relationship between building and 8.4_Glut:8.1_Glut ratio was influenced by any of the following variables: GSI, quivering, all significant RG subcluster and CDG module-related effects (RG_1_ *wdr73*, RG_2_ *wdr73*, RG_2_ CDG module score, RG_8_ CDG module score, RG_3_ *cyp19a1*, RG_4_ proportion), IEG score in all ten build-IEG+ populations, and IEG score in 8.1_Glut and 8.4_Glut (Fig. 7N). This analysis revealed RG_2_ *wdr73* expression and 4_GABA *htr1d+* IEG score as the only predicted mediators of building-associated neuronal rebalancing (Fig. 7N). To investigate candidate signals that may drive building-associated decreases in RG_2_ *wdr73* expression, we performed a similar analysis with RG_2_ *wdr73* as the outcome. This analysis revealed IEG score in 9_Glut and 4_GABA *vipr2+* as the only predicted mediators of RG_2_ *wdr73* expression (Fig. 7N). These data support a model whereby building-associated neural activity in 9_Glut and subpopulations of 4_GABA, together with RG_2_, coordinate neuronal rebalancing in Dl-v. Consistent with this model, spatial integration mapped 9_Glut to the dorsal region of the dorsal telencephalon (Dd) and 4_GABA to dorsal/supracommissural regions of the ventral telencephalon (Vd/Vs; Fig. 7C-E), both of which are reciprocally connected with Dl-v in other teleosts (O’Connell and Hofmann 2011b; Giassi, Ellis, and Maler 2012).

Among populations of interest, connection weights were weakest for RG_2_ as a receiver (Fig. 7I), consistent with a lack of direct connections between build IEG+ populations and RG_2_. In the mammalian hippocampus, neural activity can regulate glial differentiation into new neurons through “spillover”, or ligand release, diffusion, and binding to target receptors in the absence of direct synaptic connections. We reasoned that a spillover model may be sufficient to explain the data: excitation of 9_Glut, 4_GABA *vipr2+*, and 4_GABA *htr1d+* during building causes secretion of ligands that diffuse and bind to target receptors expressed on nearby RG_2_ lining the ventricular zone of Dl-v, causing RG_2_ to differentiate into new 8.4_Glut neurons. Consistent with this model, examination of ligands expressed in 9_Glut, 4_GABA *vipr2+*, and 4_GABA *htr1d+* and their paired target receptors in RG_2_ revealed NRG2-ERBB4 as the top pair (Fig. 7O). *nrg2* was one of 81 bDEGs identified in 9_Glut (Fig. 7P), and *erbb4* was preferentially expressed in RG_2_ compared to other RG, and in 8.4_Glut compared to 8.1_Glut and compared to other 8_Glut neurons (Fig. 7Q). NRG2-ERBB4 binding promotes both glial cell and neuronal differentiation and, in humans, migration of glioma cells (Ghashghaei et al. 2006; Louhivuori et al. 2018; W.-J. Zhao et al. 2021). Together our data identify a plausible circuit model whereby building-associated neural activity, together with an evolutionarily divergent gene module in glia, coordinate a cellular reorganization of Dl-v during building (Fig. 7R).

## DISCUSSION

The diversity of social behaviors in nature is an opportunity to discover how conserved genes and cell populations generate variable neural and behavioral responses to social stimuli (Johnson and Young 2018; Jourjine and Hoekstra 2021; Hofmann et al. 2014; O’Connell and Hofmann 2011a). The ability to functionally profile many heterogeneous cell populations in under- and unstudied behavioral and species systems will be a boon to this endeavor. In this study we investigated the neurobiological substrates of castle-building in *Mchenga conophoros* by integrating snRNA-seq with comparative genomics and automated behavior analysis. Using natural individual genetic variation, we matched telencephalic nuclei back to 38 test subjects, enabling powerful analyses of building-associated signals that controlled for correlated variables that may explain differences in brain gene expression. We first charted the cellular diversity of the telencephalon, and then profiled behavior- and gonadal-associated gene expression, cell type proportions, genomic signatures of behavioral evolution, and cell-cell signaling systems across telencephalic cell populations. Our results support central and related roles for glia, genome evolution, hippocampal-like neurogenesis, and cell type-specific neural activity in castle-building behavior.

### Signatures of neuronal excitation reveal candidate populations activated during building

Different social behaviors are regulated by distinct neural circuits and/or circuit activities in the brain (Newman 1999; Goodson 2005; Amadei et al. 2017; Kimchi, Xu, and Dulac 2007; Dulac, O’Connell, and Wu 2014). Identifying which cell populations are important for a specific behavior is difficult, because most tools cannot functionally profile many heterogeneous cell populations at once. Three previous studies have supported the promise of sn/scRNA-seq technologies for mapping behavior-associated IEG expression across many cell populations (Lacar et al. 2016; Moffitt et al. 2018; Y. E. Wu et al. 2017); however, all three studies were conducted in the same genetically inbred C57BL6/J mouse strain, and thus cells that were pooled for sequencing could not be matched back to individual animals. In our study, we leveraged natural genetic variation among individuals to trace ∼34,000 nuclei back to 38 individual males and analyzed building-associated IEG expression while accounting for variance explained by other biological and technical factors. Our analysis revealed novel IEG-like genes and distinct patterns of building-, quivering-, and GSI-associated neuronal excitation across clusters and gene-defined cell populations. Building was associated with increased IEG expression in 9_Glut. Spatial profiling mapped 9_Glut to Dd, a pallial region that innervates Dl in a many-to-one fashion in other fish, mirroring the conserved “pattern separator” circuit organization within the mammalian hippocampus (Elliott et al. 2017). *ntrk2+* nuclei also exhibited building-associated IEG expression, highlighting the TrkB system as a candidate player in castle-building. TrkB is a receptor that transduces activity-dependent signals into downstream modulation of neuronal differentiation, morphogenesis, survival, and long term potentiation (LTP) (Badurek et al. 2020; Lipsky and Marini 2007). Interestingly, *ntrk2+* nuclei also exhibited building-associated pNG expression, suggesting TrkB may link building-associated neuronal firing to building-associated neuronal plasticity. The only other population that exhibited both building-associated IEG and pNG expression was defined by expression of *cckbr* (encodes Cholecystokinin B Receptor). Interestingly, this receptor has recently been linked to NMDA receptor-mediated LTP and hippocampal neurogenesis in mice (Asrican et al. 2020; Chen et al. 2019).

### A role for neurogenesis in social behaviors tied to reproductive cycles

Analysis of differential gene expression, pNG expression, cluster proportions, behavior-associated genome divergence, and RG biology supported a role for neurogenesis in the evolution and expression of castle-building behavior. These analyses provided converging evidence for building-associated neurogenesis in 8_Glut, a cluster that anatomically mapped to Dl. Briefly, Dl is a brain region in the lateral pallium of fish that is thought to be homologous to the mammalian hippocampus based on gene expression, cell morphology, afferent and efferent connectivity, anatomical, and behavioral evidence (Fotowat et al. 2019; Bingman, Salas, and Rodriguez 2008; Rodríguez et al. 2002; O’Connell and Hofmann 2011b; Ganz et al. 2014; Salas et al. 2017; Ocaña et al. 2017). For example, the Dl and the hippocampus have demonstrated roles in regulating spatial learning in fish and mammals (including humans), respectively (Engelmann, Wallach, and Maler 2021; Vikbladh et al. 2019; Miller et al. 2018; Nakazawa et al. 2004). Within 8_Glut, building was associated with a shift in the relative proportions of 8.4_Glut and 8.1_Glut, two populations that mapped specifically to ventral Dl, a subregion that exhibits selective responses during spatial learning and memory formation in other fish species (Uceda et al. 2015; Ocaña et al. 2017). Our data thus support the possibility that building behavior is associated with a reorganization of hippocampal-like cell populations involved in spatial learning. Interestingly, changes in the social environment induce telencephalic cell proliferation and migration in other cichlid species within three hours, supporting the idea that behavior-associated neurogenesis can occur on relatively short timescales (Maruska, Carpenter, and Fernald 2012).

In the wild, bowers are constructed selectively during the breeding season and function as social territories that males aggressively defend against intruders, as well as mating sites for courtship and spawning with females. Bowers are constructed through thousands of spatial decisions about where to scoop and spit sand that ultimately give rise to a species-specific geometric structure. It has been reported in several species that in response to structural damage or destruction (e.g. caused by storms), males will repair or reconstruct the bower to match the size, geometry, and spatial location of the original structure (McKaye, Louda, and Stauffer 1990; Kirchshofer 1953). After the breeding season ends, bowers lose their social significance and are abandoned. Together, these data suggest that spatial learning, memory, and decision-making play a central role in bower-building, and further that spatial representations of the bower are maintained within breeding cycles. Importantly, in our paradigm, control males had previously built, suggesting that the rebalancing was temporary and eventually returned to baseline in the absence of building activity. Within this framework, it is intriguing to speculate that hippocampal-like neuronal rebalancing during building may be related to spatial representations of the bower structure and/or territory. Notably, similar phenomena have been reported in songbirds that repeat their song within a breeding season, but change their song between seasons (Brenowitz and Larson 2015; Goldman and Nottebohm 1983). These birds show robust increases in cell proliferation in vocal learning circuits during the breeding season that decline when the season is over. Neurogenesis may play an important role in seasonal mating behaviors across species, consistent with previous work demonstrating changes in brain region-specific cell proliferation and/or neurogenesis during species-specific social contexts in a variety of taxa (Walton, Pariser, and Nottebohm 2012; Bedos, Portillo, and Paredes 2018; Almli and Wilczynski 2012; Balthazart and Ball 2016; Maruska, Carpenter, and Fernald 2012; Dunlap, Chung, and Castellano 2013; Lévy et al. 2017).

### Estrogenic substrates of male social behavior

Estrogen is a female gonadal steroid hormone that can be synthesized in the male brain via conversion of testosterone to estrogen by aromatase (L. R. Nelson and Bulun 2001). In the brain, estrogen can exert its effects at multiple levels, for example by regulating gene transcription (via EREs), neuronal excitability, synaptic plasticity, neurogenesis, and G-protein coupled receptor signaling (Kelly and Rønnekleiv 2009). Multiple lines of evidence supported a potential role for estrogen in the neural coordination of building. First, bDEGs (as well as qDEGs and gDEGs) contained canonical EREs, consistent with a role for estrogen in modulating building-associated gene transcription. Out of all GO terms, ERE-containing bDEGs were most strongly enriched for “Schaffer collateral -CA1 synapse” (driven by building-associated expression of *cacng2, ppp3ca, ptprd, ptprs*, and *l1cam*), a deeply studied hippocampal synapse involved in associative learning and spatial memory in mice (Nakazawa et al. 2004; Soltesz and Losonczy 2018). In mice, estrogen increases the magnitude of long-term potentiation at this synapse (C. C. Smith, Vedder, and McMahon 2009). It is interesting to speculate that estrogen may regulate plasticity in a conserved hippocampal circuit during castle-building behavior. Second, building-associated increases in pNG expression were strongest in populations defined by neuromodulatory receptor genes, and were stronger in populations defined by ERs (*esr1, esr2, esrra, esrrb, esrrg*) compared to other receptor families, consistent with previous reports of estrogen-mediated neural plasticity in the mammalian forebrain (Barha and Galea 2010; Brinton 2009; Srivastava and Penzes 2011). Third, building was associated with strong increases in aromatase expression in RG, an effect that was driven most strongly by RG_3_. This glial population may coordinate building-associated effects of estrogen on brain gene expression, neural circuit structure and function, and male social behavior, consistent with previous work demonstrating estrogenic regulation of male social behaviors in diverse lineages (M. V. Wu et al. 2009; Huffman, O’Connell, and Hofmann 2013; Sonoko Ogawa et al. 2020; Ervin et al. 2015).

### An evolutionarily divergent gene module links neural activity and stem-like glia to hippocampal-like neurogenesis and behavior

Glial cells have recently been shown to play central roles in synaptic communication, plasticity, learning, memory, behavior, and psychiatric disease (Santello, Toni, and Volterra 2019; Kastanenka et al. 2020; Nagai et al. 2021; X. Yu et al. 2018). In addition to building-associated aromatase expression in RG, we observed building-associated changes in RG subpopulation-specific gene expression, relative proportions, and signatures of quiescence. Comparative genomic analyses across 26 behaviorally-divergent species further converged on the importance of RG in castle-building behavior, raising the possibility that transcriptional specializations in glia have served as a substrate in castle-building evolution. A module of 12 co-expressed CDGs was tightly linked to signatures of RG quiescence and was enriched in RG_2_, a population that showed building-associated downregulation of the CDG module (particularly *wdr73*), *npas3*, and markers of glial quiescence. Interestingly, one study in human epithelial cells found that suppressed *wdr73* expression was most strongly associated with increased expression of *ccnd1* (Tilley et al. 2021), an established marker of proliferation in RG/neural stem cells in vertebrates (Lukaszewicz and Anderson 2011; G. Zhang et al. 2021). Further analysis supported a circuit model whereby behavior-associated neuronal excitation of principal striatal GABAergic (4_GABA *htr1d*+ and 4_GABA *vipr2*+) and pallial glutamatergic (9_Glut) projections to 8.4_Glut nuclei in Dl-v mediate building-associated decreases in *wdr73* expression in RG_2_, which in turn mediates behavior-associated neuronal rebalancing. Examination of molecular ligand-receptor pairs expressed between build-IEG+ populations and RG_2_ suggested that a simple spillover model mediated by NRG2-ERBB4 may explain the effect. Our results thus support a model whereby castle-building evolved in part by modifying gene regulatory networks in a glial subpopulation that responds to behavior-associated neural activity and that regulates hippocampal-like neurogenesis. These data are consistent with previous work suggesting that activation of long-range projections into the hippocampus can regulate hippocampal neurogenesis (Káradóttir and Kuo 2018; Song et al. 2016).

The CDG module resides in a 19 Mbp genomic region that exhibits signals of divergence mirroring those reported for chromosomal inversions in other species systems (Lamichhaney et al. 2016; da Silva et al. 2019; Tuttle et al. 2016; Corbett-Detig and Hartl 2012; Roesti et al. 2015; Maney et al. 2020; Berg et al. 2017). It is thought that inversions can facilitate rapid evolution by protecting large-scale and adaptive cis-regulatory landscapes and multi-allele haplotypes (“supergenes”) from recombination (Schaal, Haller, and Lotterhos 2022; Hoffmann and Rieseberg 2008; Kirkpatrick and Barton 2006; Villoutreix et al. 2021). Evidence for the importance of inversions in phenotypic evolution has been shown in diverse lineages spanning flowers and humans (Huang and Rieseberg 2020; Stefansson et al. 2005). Two recent studies in the ruff and white-throated sparrows further support that inversions may shape social behavioral evolution in diverse lineages (Merritt et al. 2020; Purcell et al. 2014; Küpper et al. 2016). In our data, four genes in the CDG module, including *wdr73*, are immediately proximate to one end of the 19 Mbp region exhibiting strong behavior-associated divergence. It is therefore intriguing to speculate that these genes reside near an inversion “break point” region with a divergent cis-regulatory architecture in castle-building lineages. Future work is needed to determine if an inversion has shaped cis-regulatory expression of these genes, RG function, and the evolution of castle-building behavior in Lake Malawi cichlid fishes.

### LIMITATIONS OF THE STUDY

The molecular readout in this study was nuclear RNA which may not reflect protein function, for example due to post-transcriptional regulation. Because nuclear RNA can only be captured at a single time point within each individual, temporal analysis of decision-making making behaviors during building was limited. This study only profiled the telencephalon, and other brain regions may play critical roles in castle-building. Lastly, firing properties and circuit connections among populations can be investigated but not proven using snRNA-seq data. Future experiments are required to validate and determine the behavioral roles of specific neural circuits.

## Supporting information

S1_subject_information

S2_cluster_brain_gene_markers

S3_cluster_information

S4_cluster_marker_enrichment

S5_genes_of_interest_by_category

S6_IEG

S7_bDEG_gDEG_qDEG

S8_ERE_genes

S9_pNG_list

S10_radial_glia_quiescent_cycling_differentiation_markers

S11_CDG

S12_CDG_enrich

S13_CDG_module_enrich

S14_CDG_module_TF_correlations

S15_CellChat

## ACKNOWLEDGEMENTS

We thank our collaborators Ashley Parker and Drs. Swantje Graetsch, Manuel Stemmer, and Herwig Baier for valuable feedback during the early stages of the project; Dr. Nicholas Johnson for suggestions regarding statistical analysis of IEG co-expression; Dr. Justin Rhodes for insightful feedback on IEG expression analysis; Cristina Baker for her critical role in initial development of spatial transcriptomics wetlab pipelines; and the Georgia Tech Petit Institute Genome Analysis and Molecular Evolution Cores for their integral roles in sample processing and sequencing, respectively. This work was supported in part by NIH R01GM101095 and R01GM144560 to J.T.S., NIH F32GM128346 to Z.V.J., NIH R35 GM139594 to P.T.M., NSF Graduate Research Fellowship DGE-2039655 to T.J.L., and Human Frontiers Science Program RGP0052/2019 to J.T.S.

## CONTRIBUTIONS

### General

Z.V.J. initially conceived of the experiment and Z.V.J., J.T.S, and B.E.H. developed and designed it. Z.V.J. and B.E.H. performed all wetlab work (see details below under “Wetlab”). T.J.L. pre-processed behavioral and depth data, including in part spatial and temporal registration of both data streams and temporal anchoring to experimental endpoints. G.W.G. pre-processed snRNA-seq, DNA-seq, and spatial transcriptomics data. Z.V.J. and G.W.G. performed downstream data analysis (see details below under “Drylab”). B.E.H. matched snRNA-seq data to published neuroanatomical expression profiles (see details below under “Drylab”). Z.V.J. took the lead on writing the manuscript with critical feedback from G.W.G., J.T.S., B.E.H., and P.T.M. Z.V.J took the lead on designing and creating figures with contributions from B.E.H., G.W.G., and T.J.L., and with critical feedback from G.W.G., J.T.S., B.E.H., P.T.M., and T.J.L. J.T.S. mentored and funded Z.V.J., B.E.H., and G.W.G., and P.T.M mentored and funded T.J.L on the project. J.T.S. funded snRNA-seq, DNA-seq, and spatial transcriptomics experiments.

### Wetlab

Z.V.J. and B.E.H. developed and optimized a single nucleus isolation protocol for cichlid telencephala. Z.V.J. and B.E.H. performed all behavioral assays, surgeries, and downstream nuclei isolations for snRNA-seq. Z.V.J. performed DNA isolations for matching nuclei to subjects. B.E.H. performed all behavior assays for spatial transcriptomics. Z.V.J. and B.E.H. performed surgeries for spatial transcriptomics. B.E.H. performed all downstream wetlab work for spatial transcriptomics. The Petit Institute Genome Analysis Core at GT performed library preparation for snRNA-seq, DNA-seq, and spatial transcriptomics. The Petit Institute Molecular Evolution Core at GT performed sequencing for snRNA-seq, DNA-seq, and spatial transcriptomics.

### Drylab

Z.V.J. performed clustering and cluster marker analysis. B.E.H. systematically surveyed the literature to determine conserved neuroanatomical expression patterns of ligand, receptor, nTF, and other cell type-specific marker genes in the teleost telencephalon. B.E.H., G.W.G, and Z.V.J. collaboratively identified markers of RG quiescence, cycling, and neuronal differentiation. Z.V.J. and G.W.G. collaboratively developed many analytical approaches. Z.V.J. conducted behavioral, IEG co-expression, IEG, DEG, pNG, cell proportion, and gene set enrichment (for biological categories) analyses. G.W.G. matched nuclei to subjects and conducted comparative genomics, gene orthologue calling, ERE detection, gene module detection, and cluster enrichment (for gene lists) analyses. G.W.G. performed spatial integration of clusters and B.E.H. matched spatial transcriptomic profiles to brain regions. Z.V.J. and G.W.G. performed cell-cell communication analyses.

## STAR METHODS

### EXPERIMENTAL MODEL AND SUBJECT DETAILS

All cichlids (species *Mchenga conophoros*) used in this study were fertilized and raised into adulthood (>180 days) in the Engineered Biosystems Building cichlid aquaculture facilities at the Georgia Institute of Technology in Atlanta, GA in accordance with the Institutional Animal Care and Use Committee guidelines (IACUC protocol number A100029). This colony was originally derived from wild-caught populations collected in Lake Malawi. All experimental animals were collected as fry at approximately 14 days post-fertilization from mouthbrooding females and were raised with broodmates on a ZebTec Active Blue Stand Alone system. At approximately 60 days post-fertilization, animals were transferred to 190-L (92 cm long x 46 cm wide x 42 cm tall) glass aquaria and were housed in social communities (20-30 mixed-sex individuals) into adulthood. Environmental conditions of aquaria were similar to those of the Lake Malawi environment; subjects were maintained on a 12-h:12-h light:dark cycle with full lights on between 8am-6pm Eastern Standard Time (EST) and dim lights on for 60 minutes between light-dark transition (7am-8am and 6pm-7pm EST) in pH=8.2, 26.7°C water and fed twice daily (Spirulina Flake; Pentair Aquatic Ecosystems, Apopka, FL, U.S.A.). All tanks were maintained on a central recirculating system. Reproductive adult subject males (age 6-14 months post-fertilization, n=38) were visually identified from home tanks based on nuptial coloration and expression of classic courtship behaviors (i.e. chasing/leading, quivering). Reproductive adult stimulus females were visually identified from home tanks based on distension of the abdomen (caused by ovary growth) and/or buccal cavity (caused by mouthbrooding).

## METHOD DETAILS

### Behavior tanks

Behavior tanks were equipped with LED strip lighting synced with external room lighting to minimize large shadows and maximize consistency in video data used for action recognition (10-h:14-h light:dark cycle). Sand (Sahara Sand, 00254, Carib Sea Inc.; ACS00222) was contained within a 38.1 cm long x 45.6 cm wide section of each tank and separated from the rest of the aquarium by a custom 45.6 cm wide x 17.8 cm tall x 0.6 cm thick transparent acrylic barrier secured with plastic coated magnets (1.25 cm wide x 2.5 cm tall x 0.6 cm thick; BX084PC-BLK, K&J Magnetics, Inc.). This design ensured that all fish could freely enter and leave the enclosed sand tray region throughout the trial. At the start of the trial, the smoothed sand surface lay approximately 29.5 cm directly below a custom-designed transparent acrylic tank cover (38.1 cm long x 38.1 cm wide x 3.8 cm tall) that directly contacted the water surface to eliminate rippling for top-down depth sensing and video recordings.

### Behavior assays

Subject males were introduced to behavioral tanks containing sand and four reproductive adult age- and size-matched stimulus females of the same species. Broods were collected from all mouthbrooding females prior to introduction of subject males to behavior tanks. Prior to behavioral trials, each male was allowed to initiate castle-building to 1) confirm capacity and motivation to build and 2) minimize potential confounding effects of “novelty” on brain gene expression that may be caused by the male’s first experience building. After building was confirmed during the initial “pre-trial” period, the sand surface in each behavioral tank was smoothed shortly before lights off, and an automated depth sensing and video recording protocol was initiated as previously described using a Raspberry Pi 3 mini-computer (Raspberry Pi Foundation) (Johnson, Arrojwala, et al. 2020). Briefly, this system uses 1) a Microsoft XBox Kinect Depth sensor to track depth change across the sand surface every five minutes, enabling analysis of the developing bower structure over time, and 2) a Raspberry Pi v2 camera to record 10 hours of high-definition video data daily. The system regularly uploads depth change updates to a Google Documents spreadsheet, enabling real-time, remote monitoring of bower construction activity in each tank. Following each trial, a trained 3D Residual Network was used to predict male building and quivering behaviors from video data as previously described (Long et al. 2020).

### Tissue sampling

Actively constructing males (n=19) were identified through remote depth change updates and were collected between 11am-2pm EST (3-5 h after full lights-on and feeding) to control for potential effects of circadian rhythm, feeding, hunger, and anticipation of food on brain gene expression. At the same time, a neighboring male that was not constructing a bower (nor had initiated construction) but could also freely interact with four females and sand, was also collected (“control”, n=19). Immediately following collection, subjects were rapidly anesthetized with tricaine for rapid brain extraction, measured for standard length (SL, distance measured from snout to caudal peduncle) and body mass (BM), and rapidly decapitated for brain extraction. Telencephala (including olfactory bulbs) were dissected under a dissection microscope (Zeiss Stemi DV4 Stereo Microscope 8x -32x, 000000-1018-455), in Hibernate AB Complete nutrient medium (HAB; with 2% B27 and 0.5 mM Glutamax; BrainBits) containing 0.2 U/μl RNase Inhibitor (Sigma). Immediately following dissection telencephala were rapidly frozen on powdered dry ice and stored at -80 °C. Testes were then surgically extracted and weighed to calculate gonadosomatic index (GSI=gonad mass/BM*100) for each subject (subject information available in Table S1).

### Nuclei isolation

Nuclei were isolated following a protocol adapted from (Martelotto 2020) and optimized for cichlid telencephala. Immediately prior to single nuclei isolation, frozen telencephala were pooled into five biological replicates (n=3-4 subjects/pool) per behavioral condition (building versus control). Pools were organized such that individuals within a pool had nearly identical telencephalic mass with the aim of equalizing the relative mass of tissue and the relative number of nuclei sampled from each subject within each pool. Additionally, paired constructing versus control pools were organized such that males in both pools were matched as closely as possible in relative age, body mass, and sampling dates. Frozen telencephalon tissue sample pools were transferred into chilled lysis buffer containing 10 mM Tris-HCL (Sigma), 10 mM NaCl (Sigma), 3 mM MgCl_2_ (Sigma), 0.1% Nonidet P40 Substitute (Sigma), and Nuclease-free H_2_O. The tissue was incubated on ice and lysed for 30 minutes with gentle rotation. Following lysis, 1.0 mL HAB medium was added and the tissue was rapidly triturated 20 rounds using silanized glass Pasteur pipettes (BrainBits) with a 500 μm internal diameter to complete tissue dissociation. Dissociated tissue were centrifuged (600 rcf, 5 minutes, 4°C) and resuspended in 2.0 ml chilled wash and resuspension buffer containing 2% BSA (Sigma) and 0.2 U/μl RNase Inhibitor (Sigma, as described above “Tissue Collection”) in 1X PBS (Thermo Fisher). The nuclei suspensions were filtered through 40 μm Flowmi^®^ cell strainers (Sigma) and 30 μm MACS^®^ SmartStrainers (Milltenyi) to remove large debris and aggregations of nuclei prior to fluorescence activated cell sorting (FACS).

### Fluorescence Activated Cell Sorting

Pilot experiments revealed that multiplets (clumps of multiple nuclei adhered together) passed through both passive filtration steps, and therefore we further improved the quality and purity of our sample using FACS (BD FACSAria™ Fusion Cell Sorter, BD Biosciences). Sizing beads (6 μm; BD Biosciences) and 1.0 μg/ml DAPI (Sigma) were used to set gating parameters, enabling selection of singlet nuclei based on size (forward scatter), shape (side scatter), and DNA content (DAPI fluorescence. Thus, this step efficiently filtered out multiplets and irregularly shaped nuclei (characteristic of unhealthy or dead nuclei). At least 300,000 nuclei/pool were collected into 1 mL wash and resuspension buffer for downstream sequencing.

### snRNA-sequencing

Suspensions of isolated nuclei were loaded onto the 10x Genomics Chromium Controller (10x Genomics) at concentrations ranging from 400-500 nuclei/ul with a target range of 3,000–4,000 nuclei per sample. Downstream cDNA synthesis and library preparation using Single Cell 3’ GEM, Library and Gel Bead Kit v3.1 and Chromium i7 Multiplex Kit were performed according to manufacturer instructions (Chromium Single Cell 3’ Reagent Kits User Guide v3.1 Chemistry, 10X Genomics). Sample quality was assessed using high sensitivity DNA analysis on the Bioanalyzer 2100 system (Agilent) and libraries were quantified using a Qubit 2.0 (Invitrogen). Barcoded cDNA libraries were pooled and sequenced on the NovaSeq 6000 platform (Illumina) on a single flow cell using the 300-cycle S4 Reagent kit (2×150 bp paired-end reads; Illumina).

### DNA sequencing

Genomic DNA was isolated from diencephalic tissue sampled from each test subject using a DNeasy Blood and Tissue Kit pipeline with a 60 min lysis time and without RNase A. The 260/280 nm absorbance ratio ranged from 1.91-2.10 across subjects. Libraries were prepared following a NEBNext Ultra II FS DNA Library Prep Kit for Illumina protocol. Libraries were sequenced on two NovaSeq 6000 lanes using 300-cycle SP Reagent Kits (2×150 bp paired-end reads; Illumina).

### Spatial transcriptomics

Telencephala were microdissected from two size-matched build-control pairs of *Mchenga conophoros* males (n=4 males total), embedded in cryomolds, flash frozen on dry ice, and stored at -80°C until further processing. Tissue was cryo-sectioned coronally at 10-μm thickness at -20°C (Cryostar NX70) and mounted onto pre-chilled Visium Spatial Gene Expression slides (10X Genomics). RNA quality (RIN > 7) was confirmed using an RNA 6000 Nano Kit (Agilent). Spatial gene expression slides were processed following manufacturer instructions (Visium Spatial Gene Expression Reagent Kits User Guide, 10X Genomics). Barcoded cDNA libraries were sequenced on the NovaSeq 6000 platform (Illumina).

## QUANTIFICATION AND STATISTICAL ANALYSIS

### Behavioral Analysis

For all trials, 3D ResNet-predicted behaviors and structural change across the sand surface was analyzed over the 90 minutes preceding collection following the same general approach described previously (Johnson, Arrojwala, et al. 2020). Briefly, a smoothing algorithm was applied to remove depth change attributable to technical noise, and small regions of missing data were recovered by spatial interpolation. Bowers were defined as any region within which one-thousand or more contiguous pixels (equivalent to ∼10 cm^2^) changed in elevation by more than 0.2 cm in the same direction (∼2 cm^3^ volume change total) based on previous analysis of depth change caused by non-building home tank activity (Johnson, Arrojwala, et al. 2020). Depth change values were adjusted based on the cubed standard length of each subject male, to create a standardized measure of building activity (larger males have larger mouths and can scoop and spit a larger volume of sand). Action recognition was used to track the number, location, and timepoints of predicted bower construction behaviors (scoops, spits, and multiple events) and quivering behaviors over the same 90 min period. The number of quivering events was log-normalized due to a single outlier (building) male with 257 predicted quivering events (∼5.9 standard deviations above the mean). Feeding behaviors were not analyzed because they can be performed by both males and females and we are not able to reliably attribute individual feeding events to the subject male.

For simplicity, we generated a single “Bower Activity Index” (BAI) metric to reflect overall building activity by first calculating the regression line between depth change and building events for each trial (n=38, R^2^=0.76). We then projected each male’s depth change and bower behavior values onto that line, with the lowest value (0 predicted building events, 0 above threshold depth change) being set to 0. BAI was calculated as the Euclidean distance along the regression line from the lowest value. BAI was used as a continuous measure of castle-building behavior throughout this study.

Differences in building, quivering, and GSI between groups were analyzed using a paired t-test in which behave and control subjects collected at the same time were treated as pairs.

### nRNA-seq pre-processing and quality control

FASTQ files were processed with Cell Ranger version 3.1.0 (10X Genomics). Reads were aligned to the *Maylandia zebra* Lake Malawi cichlid genome assembly (Conte et al. 2019) using a splice-aware alignment algorithm (STAR) within Cell Ranger, and gene annotations were obtained from the same assembly (NCBI RefSeq assembly accession: GCF_000238955.4, M_zebra_UMD2a). Because nuclear RNA contains intronic sequence, they were included in the cellranger count step. Cell Ranger filtered out UMIs that were homopolymers, contained N, or contained any base with a quality score less than 10. Following these steps, Cell Ranger generated ten filtered feature-barcode matrices (one per pool) containing expression data for a total of 32,471 features (corresponding to annotated genes) and a total of 33,895 barcodes (corresponding to droplets and putative nuclei) that were used passed through additional quality control steps in the “Seurat” package in R. Examination of total transcripts, total genes, and proportion of mitochondrial transcripts were similar across all ten pools, and therefore the same criteria were used to remove potentially dead or dying nuclei from all pools. Barcodes associated with fewer than 300 total genes, fewer than 500 total transcripts, or greater than 5% (of total transcripts) mitochondrial genes were excluded from downstream analysis on this basis. This step filtered out a total of 20 (0.059%) barcodes. To reduce risk of doublets or multiplets, barcodes associated with more than 3,000 total genes or 8,000 total transcripts were also excluded. This step filtered out a total of 201 barcodes (0.59%). In total, 33,674 barcodes (99.34%) passed all quality control filters and were included in downstream analyses.

### Dimensionality reduction

In order to perform dimensionality reduction, we first identified 4,000 genes that exhibited the most variable expression patterns across nuclei using the FindVariableFeatures function in Seurat with the mean.var.plot selection method, which aims to identify variable features while controlling for the strong relationship between variability and average expression, and otherwise default parameters. Gene-level data was then scaled using the ScaleData function in Seurat with default parameters. To examine dimensionality, we first performed a linear dimensional reduction using the RunPCA command with the maximum possible number of dimensions (“dim” set to 50). We then used Seurat’s JackStraw, ScoreJackStraw, and JackStrawPlot functions to estimate and visualize the significance of the first 50 principal components (PCs), and the Elbow plot function to visualize the variance explained by the first 50 PCs. Because all 50 PCs were highly statistically significant, and no “drop off” was observed in variance explained across PCs, we used all 50 PCs for non-linear dimensional reduction (Uniform Manifold Approximation and Projection, UMAP) using the RunUMAP function in Seurat. For RunUMAP, “min.dist” was set to 0.5, “n.neighbors” was set to 50, and “metric” was set to “euclidean”.

### Clustering

Prior to clustering, nuclei were embedded into a K nearest-neighbor (KNN) graph based on euclidean distance in UMAP space, with edge weights based on local Jaccard similarity, using the FindNeighbors function in Seurat (k.param=50, dims=1:2, prune.SNN=0). Clustering was then performed using Seurat’s FindClusters function using the Louvain algorithm with multilevel refinement (algorithm=2). This final step was performed twice using two different resolution parameters to generate both coarse- and fine-grained structural descriptions of the underlying data, facilitating investigation of both major cell types as well as smaller subpopulations. For more coarse-grained clustering (resolution=0.01) identified 15 1° clusters and fine-grained clustering (resolution=1.3) identified 53 2° clusters.

### Cluster marker gene analysis

The biological identities of specific clusters were investigated using a multi-pronged approach that incorporated unbiased analysis of cluster-specific marker genes as well as supervised examination of previously established marker genes. Cluster-specific “marker” genes were identified using the FindAllMarkers function in Seurat. Briefly, this function compares gene expression within each cluster to gene expression across all other clusters and calculates Bonferroni-adjusted p-values using a Wilcoxon rank sum test. Functional enrichment analysis of GO categories among cluster-specific marker genes was investigated by first converting cichlid gene names to their human orthologs and then performing functional enrichment analysis using ToppGene Suite with default parameters. Enrichment results that survived FDR-adjustment (q<0.05) were considered statistically significant. Established cell type-specific and neuroanatomical marker genes were identified from the literature (Table S2) and were intersected with the output from FindAllMarkers to generate further insight into the biological identity of clusters.

### Assignment of nuclei to test subjects

To match individual nuclei to individual test subjects, we used Demuxlet to match variants identified in snRNA-seq reads to variants identified from genomic sequencing of each subject (Kang et al. 2018). First, genomic DNA from every test subject was collected and sequenced. In total, 276.7 Gbp of sequenced reads were assigned quality scores≥30 (91.4% of all reads). The corresponding FASTQ files were filtered and aligned to the M. zebra Lake Malawi cichlid genome umd2a assembly (NCBI RefSeq assembly accession: GCF_000238955.4, M_zebra_UMD2a). The resulting bam file was sorted, duplicates removed, read groups added, and indexed using Picard tools. Variants were then called using GATK v4.1.8.1 HaplotypeCaller using the M. zebra umd2a reference genome. Based on pool, individual vcf files were merged, resulting in 10 files (one for each pool). These files were then filtered to retain only variants that varied among individuals in a pool. For each pool, only SNPs for which 1) at least one individual from the pool had a different genotype from the other individuals, and 2) no individuals had missing data, were used as input to Demuxlet. The number of SNPs used ranged from 112,385 to 357,177 with a mean of 241,780±22,369 per pool.

Next, variants were called from snRNA-seq reads following a similar pipeline. Reads from Cell Ranger’s output bam file were filtered for those that passed the quality control metrics described above using samtools v1.11. The resulting bam file was sorted, duplicates removed, read groups added, and indexed using Picard tools. Variants were then called using GATK HaplotypeCaller using the M. zebra umd2a reference genome and without the MappingQualityAvailableReadFilter to retain reads that were confidently mapped by Cell Ranger (MAPQ score of 255). The SNPs from the snRNA-seq reads and the genomic DNA were input to Demuxlet, which computed a likelihood estimation that each nucleus belongs to each individual in the pool. Nuclei were assigned to the individual test subject with the greatest probability estimated by Demuxlet.

### Identification of IEG-like genes

Three canonical IEGs (*c-fos, egr1, npas4*) were used to identify additional genes exhibited IEG-like expression across clusters. For each of these three IEGs, nuclei were split into IEG-positive versus IEG-negative nuclei within each of the 53 clusters. Within each cluster, differential gene expression was analyzed between IEG-positive versus IEG-negative nuclei using the FindMarkers function in Seurat, with “logfc.threshold” set to 0, and “min.pct” set to 1/57 (57 was selected as this was the number of nuclei in the smallest cluster). Within each cluster, any genes that did not meet this criterion were excluded and were assigned a p-value of 1. Because FindMarkers requires at least three nuclei to be present in both comparison groups, clusters that contained less than three IEG-positive nuclei were excluded. Genes that were detected in the majority of clusters, and that were significantly (p<0.05) upregulated in IEG-positive nuclei in the majority of those clusters were considered to be significantly co-expressed with each individual IEG. Genes that were significantly co-expressed with all three IEGs were used as IEG-like markers for downstream analyses of IEG-like expression.

### Differential IEG expression

Building-, quivering-, and gonadal-associated IEG expression was analyzed in 1° and 2° clusters, gene-defined populations within 1° and 2° clusters, and gene-defined populations regardless of cluster. To do this, we calculated an IEG score for each nucleus, equal to the number of unique IEG-like genes (n=25) expressed. Building-, quivering-, and gonadal-associated differences in IEG score were analyzed using a beta-binomial model in which the number of IEG-like genes observed as well as the number of the IEG-like genes not observed were tracked as indicators of recent neuronal excitation. This analysis was performed using the ‘BBmm’ package in R (m=25). Because castle-building, quivering, and GSI were correlated with one another, we analyzed expression using a sequence of beta-binomial mixed-effects models in which different pairwise combinations of predictor variables (building, quivering, and GSI) competed to explain variance in IEG score. These models also included nested random terms to account for variance explained by individual variation, pair, pool, and RNA isolation/cDNA library generation batch. Within this framework, we ran the following seven models, which allowed building (analyzed as either a binary or a continuous variable), quivering, and GSI to compete in all possible combinations to explain variance in IEG score:

1. IEG score ∼ **BAI** + **log(quivering events)** + *(subject/pool/batch) + (subject/pair)*
2. IEG score ∼ **BAI** + **GSI** + *(subject/pool/batch) + (subject/pair)*
3. IEG score ∼ **BAI** + **log(quivering events)** + **GSI** + *(subject/pool/batch) + (subject/pair)*
4. IEG score ∼ **Condition** + **log(quivering events)** + *(subject/pool/batch) + (subject/pair)*
5. IEG score ∼ **Condition** + **GSI** + *(subject/pool/batch) + (subject/pair)*
6. IEG score ∼ **Condition** + **log(quivering events)** + **GSI** + *(subject/pool/batch) + (subject/pair)*
7. IEG score ∼ **log(quivering events)** + **GSI** + *(subject/pool/batch) + (subject/pair)*

We defined significant building-, quivering-, and gonadal-associated IEG effects as those in which 1) the raw p-value for the corresponding fixed effect (for building, BAI and condition; for quivering, log-normalized quivering; for gonadal, GSI) was significant (p<0.05) in every model, and 2) the harmonic mean p-value across models was significant after adjusting for multiple comparisons for all genes and populations analyzed (hmp_adj_<0.05). To calculate the harmonic mean p-value, we used the “harmonicmeanp” package in R. Thus, building-associated IEG effects were significant (the raw p-value for the effect of “condition” and “BAI” <0.05) in models 1-6, and if the harmonic mean p-value across models 1-6 was significant after adjusting for multiple comparisons across all cell populations.

### Building-, quivering-, and gonadal-associated gene expression

Building-, quivering-, and gonadal-associated gene expression was analyzed within 1° and 2° clusters using a multiple linear mixed-effects regression approach with the “glmmSeq” package in R (https://github.com/KatrionaGoldmann/glmmSeq). Because castle-building, quivering, and GSI were correlated with one another, we analyzed expression using a sequence of linear mixed-effects regression models in which different pairwise combinations of predictor variables (building, quivering, and GSI) competed to explain variance in gene expression. These models also included nested random terms to account for variance explained by individual variation, pair, sample pool, and 10X Chromium run. Thus, sample size was equal to the number of individuals (n=38), with many repeated observations being recorded from each individual (equal to the number of nuclei sampled from that individual as assigned to the cluster being analyzed). Building was analyzed both as a continuous variable (BAI) and as a binary categorical variable (behave versus control).

We defined bDEGS, qDEGs, and gDEGs as genes (within clusters) in which expression was significantly (raw p<0.05) associated with the corresponding fixed effect (for bDEGs, BAI and condition; for qDEGs, log-normalized quivering; for gDEGs, GSI) in every model, and additionally in which the harmonic mean p-value across models was significant after adjusting for multiple comparisons for all genes and all clusters (5% false discovery rate). For each model, dispersion was estimated for each gene using the “DESeq2” package in R, using parameters recommended for single cell datasets (fitType = “glmGamPoi”, minmu = 1e-06). Size factors for each gene were calculated using the “scran” package in R, using default parameters, except that max.cluster.size was set to the number of nuclei assigned to the cluster being analyzed. Genes that were not observed in 19/19 pairs were excluded from analysis.

### Estrogen response element detection

Estrogen receptors are hormone-dependent transcription factors capable of regulating target gene expression by binding to specific DNA sequences called estrogen response elements (EREs). EREs can be easily identified by their prototypic motif of AGGTCA separated by a 3-base spacer (Ikeda, Horie-Inoue, and Inoue 2015). Genes with an ERE motif less than 25 kilobases away were found and the location of the ERE relative to the gene was recorded as either intragenic (ERE within the start site to the 3’ polyA tail), promoter (ERE <= 5 kb upstream of the gene), or distal (all other locations less than 25 kb away from the gene). To identify the location of the ERE to the closest gene, bedtools v2.29.1 was used using the closest command.

### Building-, quivering-, and gonadal-associated pNG expression

Building-, quivering-, and gonadal-associated pNG expression was analyzed in 1° and 2° clusters, gene-defined populations within 1° and 2° clusters, and gene-defined populations regardless of cluster using the same general approach described for IEG expression, except that building-associated effects were defined as those that were significantly associated with condition in all models. Because we did not expect neurogenesis or associated cellular processes to proceed over 90-minute timescales, we did not additionally require effects to be significantly associated with BAI in all models.

### Building-associated changes in cell proportions

Behavior-associated differences in cell type-specific proportions were analyzed for 1° and 2° clusters with a binomial mixed-effects regression model using the glmer function within the “lmer” package in R. The model included condition, GSI, and quivering as fixed effects, and included a random term for individual variation. 1° cluster proportions were calculated as the proportion of all nuclei assigned to each 1° cluster, and 2° cluster proportions were calculated as the proportion of 1° “parent” cluster nuclei assigned to each 2° “daughter” cluster. Thus, each nucleus was treated as an observation with a binary outcome (either an instance of the target cluster or not) from an individual that could be explained by condition, quivering, or GSI. p-values were estimated using the ‘lmerTest’ package in R, and qvalues were calculated using the ‘qvalue’ package in R. Building-associated effects were defined as those that were significant after accounting for multiple comparisons across all clusters with a false discovery rate of 5% (q<0.05).

### Cluster-specific enrichment of gene sets

To test if genes associated with the evolution of bower construction behavior (identified through comparative genomics) were enriched in specific cell populations, we first calculated a “gene set score” for each nucleus, equal to the total number of unique behavioral evolution genes expressed. Because the gene set score could be impacted by the total volume of sequence data sampled from each nucleus, we divided the gene set score by the total number of unique genes expressed in each nucleus. To quantify enrichment, a Z-test was then used to compare “normalized” gene set scores for all nuclei within a cluster compared to all other nuclei. The distribution of the normalized values was assumed to be normal according to the central limit theorem and population standard deviation was approximated using sample standard deviation.

Secondly, the effect size, as measured by Cohen’s d, of the results were compared to those of random gene lists. To prevent differences in overall amount of expression between random genes and genes of interest from skewing results, random genes lists were chosen that had approximately equal number of UMIs expressed as a whole to the genes of interest. This was achieved by first ranking all the genes from the highest number of UMIs expressed to the lowest. Next, for each gene of interest, a pool of 100 random genes were chosen that were ranked most closely to the gene of interest and were not a gene of interest themselves. Then, 10,000 random gene lists were created by choosing one gene at random from each pool. The enrichment test described above was then performed on the random gene lists. Finally, clusters that were significantly enriched for the genes of interest according to the process above and had significantly greater effect sizes than the 10,000 random gene lists were considered to be significant.

### RG subclustering

RG subclusters were determined using the same general procedure used for clustering 1° and 2° clusters but restricted to only those nuclei assigned to 1.1_RG and 1.2_RG.

### Analysis of castle-associated genomic divergence

In order to identify potential behavior-associated genomic variants, comparative genomic analyses were performed using genomic sequence data collected from 27 Lake Malawi cichlid species (Patil et al. 2021). Fixation indices (F_ST_) were calculated for polymorphic variants in two separate analyses using vcftools v0.1.17. The first analysis compared pit-digging (N=11) versus castle-building (N=9) species, and the second compared rock-dwelling (N=7) versus castle-building (N=9) species. Variants for which sequence data was missing from 50% or more of species in either group were excluded from analysis. F_ST_ analyses were performed separately using the --weir-fst-pop and --fst-window-size 10000 flag to calculate F_ST_ across 10kb bins in vcftools. Then, bins where F_ST_ was greater than 0.20 in the pit-castle comparison and 0.20 in the rock-castle comparison were kept. These thresholds are both greater than the minimum F_ST_ of FDR-adjusted significant bins. By creating these more strict thresholds we aimed to ensure that the selected bins were extremely divergent between castle-building and non-castle-building species. Additionally, a higher threshold was selected for the rock-castle than the pit-castle comparison because of the greater evolutionary distance and thus greater overall F_ST_. Finally, genes that were within 25kb of these bins meeting these thresholds were identified using bcftools v1.11 with the closest command and the *M. zebra* genome as reference. Genes within 25kb of highly divergent pit-castle and rock-castle bins are referred to here as “castle-divergent”.

### CDG co-expression and module analysis

Modules of co-expressed CDGs were analyzed using weighted correlation network analysis (WGCNA) using the “WGCNA” v1.70-3 package in R. CDGs that were not observed in any nucleus were excluded from analysis. The normalized gene expression data for CDGs was used as the input gene expression matrix and the function pickSoftThreshold was used to determine the optimal soft-thresholding power. We determined the optimal soft-thresholding power to be 1 because it was the lowest power for which the scale-free topology fit index reached 0.90. Then an adjacency matrix was created from the input gene expression matrix using the adjacency function with power = 1, type = “signed” and otherwise default parameters. The adjacency matrix was used as the topological overlap matrix (TOM) and the dissimilarity matrix was calculated as 1 – TOM. To detect modules, k-means clustering was performed using all possible values of k and the results were compared to determine the optimal k. First, a distance matrix was constructed from the dissimilarity matrix produced by WGNCA using the dist function in R. Next, the function pam from the R package “cluster” v2.1.0 was used to cluster the distance matrix with diss = T, otherwise default parameters, and k set to the value that produced the highest average silhouette width for all genes. Briefly, silhouette width is a measure of the similarity of the genes within a module to the genes outside the module, and higher values indicate better clustering. We found that k=2, had the greatest average silhouette width. The strength of the module was evaluated using a two-sampled Welch t-test comparing the silhouette width and gene-gene correlations for CDGs within the module versus CDGs outside the module. To analyze the relationship between the CDG module and signatures of RG quiescence, the correlation coefficient was calculated based the number of genes in the CDG module expressed in each nucleus versus the number of quiescent markers expressed in each nucleus. We compared the correlation coefficient against a permuted null distribution by randomly shuffling the expression values of each gene in the module 10,000 times.

### Spatial transcriptomic pre-processing and quality control

Base Call files were demultiplexed into FASTQ files and processed with Space Ranger v1.3.1 (10X Genomics). Reads were aligned to the *M. zebra* umd2a reference assembly as described above for snRNA-seq (Conte et al. 2019). Following these steps, Space Ranger generated three filtered feature-barcode matrices containing expression data for a total of 32,471 features (corresponding to annotated genes). Spots with 0 UMIs were removed resulting in 6,707 spots used in downstream analysis.

### Spatial integration of snRNA-seq clusters

To predict locations of specific snRNA-seq identified clusters in spatial transcriptomics data, an ‘anchor’-based integration workflow in Seurat was used. First, both the snRNA-seq and spatial data were normalized using the SCTransform in Seurat. Next, anchors were identified between the reference snRNA-seq and the query spatial data using FindTransferAnchors in Seurat, and a matrix of predictions scores was generated for each cluster in every spot using the TransferData function in Seurat. The maximum prediction score across clusters was not uniform, therefore we normalized the values between 0 and 1 in order to enable meaningful comparisons across cell types.

### Cell-cell communication analysis

To assess the connectivity of cell populations, cell-cell communication analysis was performed using the R package CellChat v1.5.0. Briefly, CellChat estimates the strength of potential connection between populations from a gene expression matrix based on a database of human ligand-receptor interactions. In order to find the connection strengths between primary and secondary clusters, the gene expression matrix was duplicated and the cells in the first copy were assigned the primary labels and the cells in the second copy were assigned secondary labels. We also sought to analyze connections among additional gene-defined populations that demonstrated behavior-associated IEG expression. To achieve this, the gene expression matrices for cells from these populations were duplicated againTherefore, before connection strengths were evaluated, the human orthologs of the *M*.*zebra* gene names in the gene expression matrix were found. Since, the gene expression matrix does not allow for duplicate gene names, e.g. many-to-one orthologs, values for the many-to-one ortholog with the greatest number of normalized counts were kept and others were excluded from analysis. Next, a CellChat object was created using the createCellChat function.

Over-expressed genes and over-expressed interactions were found using the identifyOverExpressedGenes and identifyOverExpressedInteractions functions respectively. Next, connection strengths were calculated using the computeCommunProb function with the method for computing the average gene expression per cell group set to truncatedMean, trim set to 0.1, and population.size set to FALSE. Then, the cellular communication network was inferred and aggregated using the filterCommunication and aggregateNet functions. The receptor-ligand and the signalling pathway weights were saved using subsetCommunication with the slot.name parameter set to ‘net’ and netP respectively.

### Regularized Multiple Mediation Analysis

Regularized multiple mediation analysis was performed using the R package mma v10.6-1(Q. Yu and Li 2017). Briefly, this analysis used a regularization approach to test if one or more mediators explained the relationship between building and 8.4_Glut:8.1_Glut neuronal rebalancing, and between building and RG_2_ *wdr73* expression. The following variables were considered as possible mediators of building-associated 8.4_Glut:8.1_Glut neuronal rebalancing: GSI, quivering, RG_1_ *wdr73* expression, RG_2_ *wdr73* expression, RG_2_ CDG module score, RG_8_ CDG module score, RG_3_ *cyp19a1* expression, RG_4_ proportion, IEG score in all ten build-IEG+ populations, and IEG score in 8.1_Glut and 8.4_Glut. To investigate possible mediators of RG_2_ *wdr73* expression, the same variables were analyzed, except RG_2_ *wdr73* expression and RG_2_ CDG module (which contains *wdr73*) score were excluded as possible mediators. The analysis was performed with testtype = 1 (LASSO) and alpha1 and alpha2 both set to 0.05.

